# Stress-dependent inhibition of cell polarity through unbalancing the GEF/GAP regulation of Cdc42

**DOI:** 10.1101/2021.08.10.455852

**Authors:** Clàudia Salat-Canela, Mercè Carmona, Rebeca Martín-García, Pilar Pérez, José Ayté, Elena Hidalgo

**Author notes:** Correspondence should be addressed to J.A., P.P. or E.H.;. Lead contact: E.H.

## Abstract

Cdc42 rules cell polarity and growth in fission yeast. It is negatively and positively regulated by GTPase-activating proteins (GAPs) and by Guanine nucleotide Exchange factors (GEFs), respectively. Active Cdc42-GTP localizes to the poles, where it associates with numerous proteins constituting the polarity module. However, little is known about its down-regulation. We describe here that oxidative stress causes Sty1 kinase-dependent Cdc42 inactivation at cell poles. Both the amount of active Cdc42 at poles and cell length inversely correlate with Sty1 activity, explaining the elongated morphology of *Δsty1* cells. We have created stress-blinded cell poles by either eliminating two Cdc42 GAPs or through the constitutive tethering of a GEF to the cell tips, and biochemically demonstrate that Rga3 is a direct substrate of Sty1. We propose that stress-activated Sty1 promotes GTP hydrolysis and prevents GEF activity at the cell tips, thus leading to the inhibition of Cdc42 and polarized growth cessation.

## INTRODUCTION

Ras-homologous (Rho) GTPases regulate cell polarity and growth in eukaryotes. In particular, Cdc42 has a central role in the establishment of polarized growth. In budding yeast, Cdc42 localizes exclusively at the site of bud growth during division, but this model system cannot be used to explain multiple growing areas such as the ones existing in other cell types including fission yeast. *Schizosaccharomyces pombe* cells not only display Cdc42 activity sites at the area of cell division, when an actomyosin ring assembles at the center of the mother cell and contracts to give rise to two daughter cells, but it also display monopolar Cdc42-dependent growth from the old end just after cell division, and shifts to a bipolar mode when it reaches a certain length in a process known as NETO (new end take off) (Mitchison and Nurse, 1985). Therefore, *S. pombe* is an excellent model to study multiple polarity sites, and the shape of fission yeast is largely dependent on the Cdc42 polarity module. Thus, wild-type cells have a rod-like phenotype, while decreased Cdc42 levels lead to a rounded phenotype (Miller and Johnson, 1994).

Cdc42 localization and activation are tightly regulated in space and in time. As with all G-proteins, Cdc42 acts as a switch that can be found in a GTP- or a GDP-bound state, and this modulates its ability to interact with other components of the polarity module. Therefore, different GEFs (guanine-nucleotide-exchange factors) activate Cdc42 and GAPs (GTPase-activating proteins) negatively regulate its activity. Two GEFs have been characterized for Cdc42 in fission yeast: Scd1, which localizes at cell tips with GTP-Cdc42 during interphase (Kelly and Nurse, 2011), and Gef1, with apparent cytoplasmic localization (Tay et al., 2018). Both GEFs are found at the cytokinetic ring during cell division (Coll et al., 2003; Hirota et al., 2003). Regarding GAPs, a negative regulator GAP was early described for Cdc42, Rga4, which localizes to the cell sides and blocks the spreading of active Cdc42 out of cell tips (Das et al., 2007; Tatebe et al., 2008). Similar lateral localization has been reported for Rga6, which has been proposed to collaborate with Rga4 to spatially restrict active Cdc42 to cell tips (Revilla-Guarinos et al., 2016). Rga3 has been recently described as a Cdc42 GAP localized to cell tips, but its absence does not lead to any detectable phenotype during mitotic cell growth. Instead, Rga3 does play a role during sexual reproduction by regulating Cdc42 activity at the polarity patch (Gallo Castro and Martin, 2018).

Most Rho GTPases, including Cdc42, are localized to the cell membrane thanks to the prenylation of a C-terminal cysteine residue within a CAAX box (Choy et al., 1999); thus, a third family of regulators, GDIs (GDP-dissociation inhibitors) interact with Rho GTPases and mask their modified C-terminal domain, what disturbs its interaction with the lipid bilayer and maintains the Rho protein at the cytosol (DerMardirossian and Bokoch, 2005). In contrast to other yeast, *S. pombe* Cdc42 local activation does not rely on GDI-mediated extraction (Bendezu et al., 2015). Other factors such as the concentration of this small GTPase at the plasma membrane are also tightly regulated during the cell cycle, and in general the activity of Cdc42 (GTP vs. GDP forms) affects its membrane localization and steady-state levels (Estravis et al., 2017).

How are new polarity sites established at a given position? Both in *Saccharomyces cerevisiae* and *S. pombe*, the formation of a quaternary complex between Cdc42, a p21-activated kinase (PAK1/CLA4 or Pak1), a scaffolding protein (BEM1 or Scd2) and the GEF (CDC24 or Scd1) is crucial to enhance local activation of Cdc42. The recruitment of the GEF for GTP-Cdc42 at the membrane favors the activation of neighbor Cdc42 molecules thus creating a positive feedback loop that contributes to the creation of new polarity sites [for a review, see (Chiou et al., 2017)]. Moreover, in fission yeast the distinct spatial distribution of GEFs and GAPs is also vital to seed sites of polarity. Therefore, the landmark established by Tea1 and Tea4, driven to the cell tips by microtubules, promotes local activation of Cdc42 at cell tips by excluding Rga4 (Kokkoris et al., 2014). Once GTP-Cdc42 is localized to growth sites, the polarity module organizes cytoskeletal elements through a collection of effectors, such as formins, to yield the specific polarized morphology.

While positive feedback loops probably explain localization in space and time at one particular position of the Cdc42 module in either yeasts [for reviews, see (Chiou et al., 2017; Martin, 2015)], negative Cdc42 regulators are required to explain the oscillatory behavior of the module in post-NETO bipolar fission yeast cells (Das et al., 2012), and to limit the spreading of the polarity cluster. A relevant example of the later is the inhibitory phosphorylation of *S. cerevisiae* CDC24, the main GEF of CDC42, by the downstream effector PAK, to downregulate polarity sites as a negative feedback loop (Gulli et al., 2000; Kuo et al., 2014; Wai et al., 2009).

Stress signals mediated by the MAP kinase Sty1 pathway in fission yeast cause dispersion of Cdc42 (Mutavchiev et al., 2016). Thus, latrunculin A (LatA)-activated Sty1 promotes GTP-Cdc42 dispersal from cell tips. In response to environmental stresses, Sty1 gets activated and triggers a transcriptional core environmental stress response, mainly based on the transcription factor Atf1. Thus, over 400 genes are up-regulated more than two-fold after hydrogen peroxide (H_2_O_2_) and other environmental stresses which threaten yeast viability. Stress-dependent phosphorylation of the MAP kinase, which is kept inactive in the absence of stress by several tyrosine and serine/threonine phosphatases (such as Pyp1,2 and Ptc1/4, respectively) (Millar et al., 1995; Shiozaki and Russell, 1995b), triggers Sty1 nuclear accumulation and phosphorylation of Atf1 and, probably, other transcriptional mediators (Salat-Canela et al., 2017; Wilkinson et al., 1996). Cells lacking Sty1 show reduced tolerance to many environmental stresses such as heat shock, oxidative stress and nutritional deprivation (Toone et al., 1998; Zuin et al., 2010). These *Δsty1* cells display an elongated ‘*cdc*’-like phenotype, suggesting that Sty1 may also have a role during the G2/M cell cycle transition under non-stressed conditions (Shiozaki and Russell, 1995a).

Here, we have investigated whether environmental stress regulating the Sty1 pathway could modulate Cdc42 activity at the growth area, and whether this regulation could explain the elongated phenotype of cells lacking Sty1. We show here that H_2_O_2_, which transiently activates Sty1, also triggers active Cdc42 dispersal from the cell tips in a Sty1-dependent manner without promoting actin depolymerization, and stress-independent Sty1 activation in a *Δpyp1* background is sufficient to induce these effects on polarity. We demonstrate that Sty1 activity inversely correlates with active Cdc42 at cell tips and, importantly, with cell length. Furthermore, Sty1 inhibitory role on Cdc42 activity may be exerted through activation at cell tips of Rga3 and Rga6 GAPs, and the combined Gef1 delocalization and Scd1 inactivation, with a net transient inhibition of Cdc42 and growth cessation.

## RESULTS

### H_2_O_2_triggers a Sty1-dependent and actin depolymerization-independent inactivation of Cdc42 at cell tips

In the presence of the actin depolymerization drug LatA, the active Cdc42 polarity module is dispersed from cell tips in a Sty1-dependent manner (Mutavchiev et al., 2016). The MAP kinase Sty1 is strongly activated by environmental cues, including oxidative stress, to trigger a massive gene expression program (Chen et al., 2003). We used GFP-tagged CRIB (Cdc42/Rac interactive binding motif) to monitor the presence of active, GTP-bound Cdc42 in the presence of Sty1-activating doses of H_2_O_2_ (Ozbudak et al., 2005; Tatebe et al., 2008). As shown in Fig. 1A, treatment of wild-type cells with either LatA or H_2_O_2_ causes dispersal of active Cdc42 from the cell tips to lateral patches (indicated with arrows). The loss of CRIB-GFP at cell poles follows similar kinetics upon LatA and H_2_O_2_ treatments (Fig. 1B). We used Lifeact-mCherry as a marker to label F-actin in living cells (Huang et al., 2012). As reported before (Mutavchiev et al., 2016), actin depolymerization upon LatA imposition is faster than CRIB-3GFP dispersal from cell tips (Fig. 1A), which suggests that it does not drive Cdc42 inactivation at poles. Indeed, H_2_O_2_ treatment does not significantly affect actin cables (Fig. 1A), and nevertheless causes CRIB-3GFP dispersal. Both types of treatments, the irreversible inhibitor LatA and the transient, natural stressor H_2_O_2_ cause a permanent or temporal growth inhibition, respectively, as demonstrated with growth curves (Fig. S1A) and measuring net cell elongation rates after stress imposition in time-lapse experiments (Fig. 1C and Fig. S1B). In conclusion, oxidative stress causes inactivation of the polarity module at cell tips, which is concomitant to cell growth cessation.

**Figure 1.**
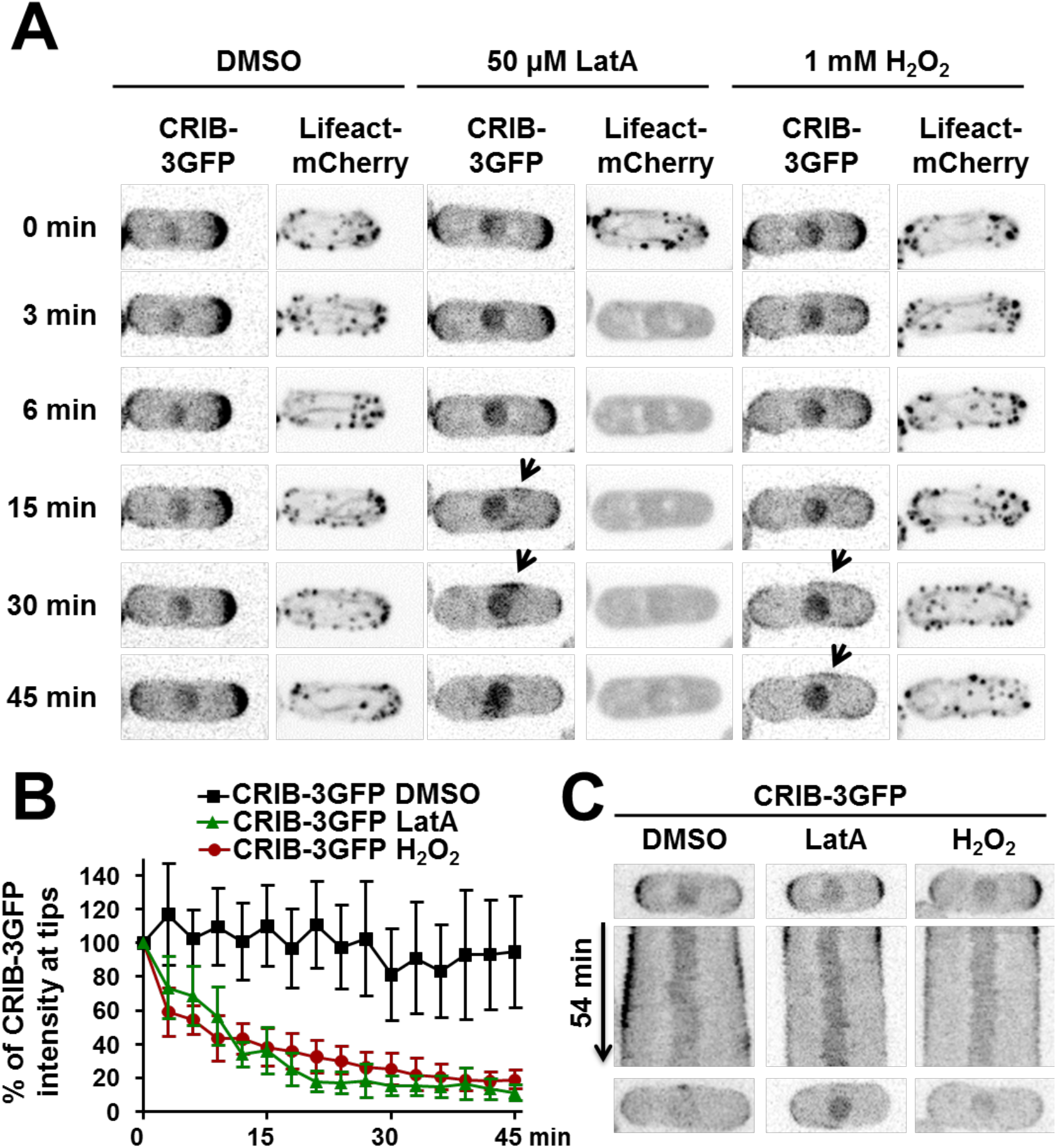
Activation of Sty1 MAP kinase by oxidative stress leads to Cdc42 depolarization from cell tips. **(A)** H_2_O_2_ imposition causes GTP-Cdc42 depolarization from cell tips at similar kinetics than LatA treatment. Still images showing GTP-bound Cdc42 and F-actin staining. Several points from time-lapse experiments of YE cultures of strain CS208 (*CRIB-3GFP Lifeact-mCherry*) during DMSO, 1 mM H_2_O_2_ or 50 µM LatA treatments are shown. Arrows indicate the formation of Cdc42-GTP lateral patches. **(B)** Quantification of CRIB-GFP intensity at cell tips from time-lapse experiments as in A. Mean, standard deviation (SD) are shown. At least 10 different cells were analysed for each condition. **(C)** Cell growth ceases after treatment with LatA or H_2_O_2_. Kymographs showing CRIB-3GFP cells from time-lapse experiments as in A.

### Stress-dependent positioning of the Cdc42 polarity module to lateral patches is dependent on the Cdc42 GEF Gef1

As shown above for H_2_O_2_ and before for LatA treatments (Bendezu and Martin, 2011; Mutavchiev et al., 2016), stress imposition causes CRIB dispersal from the tips to lateral patches close to the cell center (see arrows in Fig. 1A). We tested the presence of different Cdc42 module components at both cell sites before and after stress imposition. As shown in Fig. 2A, the main Cdc42 GEF Scd1 and its scaffold Scd2 are localized to cell tips prior to stress, and are totally (Scd1-GFP) or partially (Scd2-GFP) dispersed from the poles upon peroxide treatment. The downstream effector kinase Pak1 is also dispersed from the poles after stress imposition, probably as a consequence of Cdc42 inactivation (Fig. 2A). Upon exit from the cell tips, Scd2 and Pak1, but not Scd1, are present at the lateral patches of active Cdc42 (see white arrows in Fig. 2A), suggesting that a GEF different to Scd1 is responsible for the active lateral surfaces which appear after stress imposition. In fact, the Scd1 scaffold, Scd2, is not required for the formation of the patches (Fig. S2A). As shown in Fig. 2A, Gef1-3YFP is located at the cytosol but also decorates the tips of non-dividing cells prior to stress; the localization at tips is lost upon H_2_O_2_ stress, with Gef1-3YFP clearly localizing to lateral patches 45 min after stress imposition (see white arrow). Indeed, while dispersal of CRIB-3GFP from cell tips upon H_2_O_2_ (Fig. 2B) or LatA (Fig. S2B) treatments is not blocked in strain *Δgef1*, the formation of the lateral active patches is completely dependent on the presence of Gef1 (Fig. 2B and Fig. S2B).

**Figure 2.**
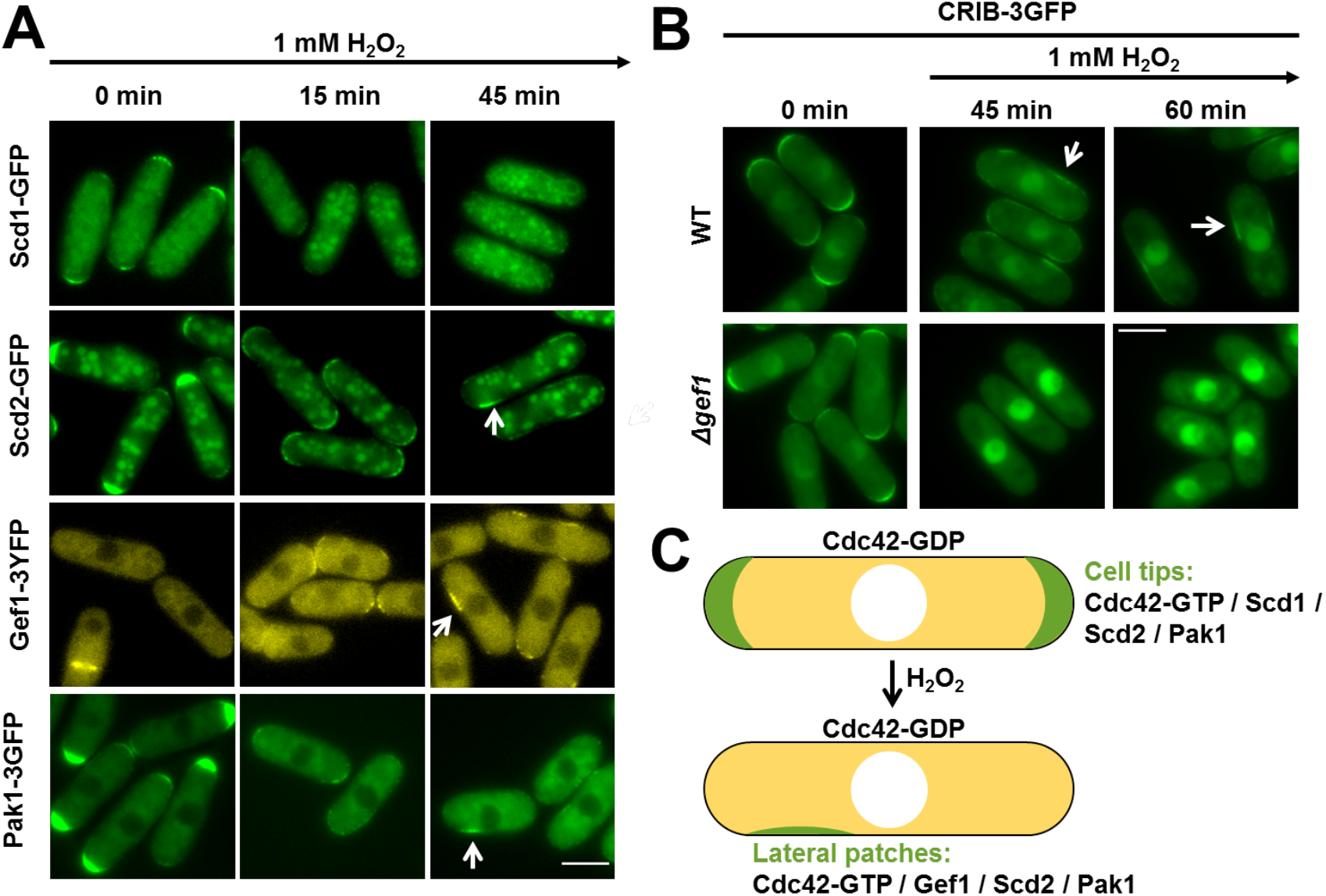
GTP-Cdc42 lateral patches are dependent on Gef1. **(A)** Scd2 and Pak1 proteins, together with Gef1, but not Scd1, are localized in Cdc42-GTP lateral patches after stress imposition. Representative images of GTP-bound Cdc42 from logarithmic YE cultures of strains PPG56.66 (*scd1-GFP*), PPG142.42 (*scd2-GFP*), PPG69.03 (*pak1-3GFP*) and FV1218 (*gef1-3YFP*) treated with 1 mM H_2_O_2_ for the indicated time-points. White arrows indicate the presence of Cdc42-GTP lateral patches. Scale bar: 5 µm. **(B)** Activation of Cdc42 at cortical regions exclusively depends on Gef1. Representative images of GTP-bound Cdc42 from logarithmic YE cultures of strains PPG65.60 (*CRIB-3GFP*) and PPG70.07 (*CRIB-3GFP Δgef1)* treated with 1 mM H_2_O_2_ for the indicated time-points. White arrows indicate the presence of Cdc42-GTP lateral patches. Scale bar: 5 µm. **(C)** Scheme depicting the localization of some Cdc42 polarity module components before and after stress imposition. During interphase, Cdc42 is active at cell tips, where it forms a quaternary complex with the GEF Scd1, the scaffold Scd2 and the effector kinase Pak1. After H_2_O_2_ treatment, GTP-Cdc42 is dispersed from cell tips and it is localized at active lateral patches together with the GEF Gef1, Scd2 and Pak1.

In parallel to our studies, other research groups obtained similar evidences using different stressors (Chen et al., 2019; Hercyk et al., 2019). We conclude that stress promotes redistribution of the active Cdc42 polarity module from the cell tips towards side patches, which contain and require Gef1 but not Scd1 (Fig. 2C). We will use hereafter these lateral sites as a secondary hallmark of stress-dependent cell growth arrest.

### Active Sty1 directly promotes the inactivation of Cdc42 at cell tips

As reported above, oxidative stress can cause inactivation of Cdc42 at cell poles, cell growth arrest and the appearance of Gef1-dependent lateral patches of active Cdc42. To confirm that active Sty1 directly drives the redistribution of the active Cdc42 polarity module from the poles to the sides of the cell, we have pursued three types of strategies: (i) we have first tested the effect of H_2_O_2_ imposition in CRIB-3GFP dispersal in cells lacking Sty1; (ii) we have then analyzed whether activation of the Sty1- and Atf1-dependent anti-stress gene expression program is required to trigger H_2_O_2_-dependent cellular depolarization by expressing CRIB-3GFP in cells lacking Atf1; and (iii) we have studied the effect on active Cdc42 in genetically engineered cells expressing a signal-independent active Sty1 allele.

We expressed CRIB-tdTomato (Revilla-Guarinos et al., 2016) in wild-type and *Δsty1* cells overexpressing catalase, to minimize the direct toxic effect of intracellular peroxides in cells lacking the MAP kinase. Over-expression of catalase in wild-type cells does not alter the inhibitory effect of stress on active Cdc42 at cell tips (Fig. 3A and Fig. S3A). We demonstrate that active Cdc42 is not redistributed from cell poles to cell sides upon H_2_O_2_ addition in cells lacking the MAP kinase (Fig. 3AB), and cell growth is not inhibited (Fig. 3C). Secondly, the absence of the transcription factor Atf1, required for the activation of the massive gene expression program (Chen et al., 2003; Chen et al., 2008), does not largely affect CRIB-3GFP dispersal from cell tips upon stress imposition (Fig. S3BC), which suggests that Sty1 kinase activity is itself required at cell tips.

**Figure 3.**
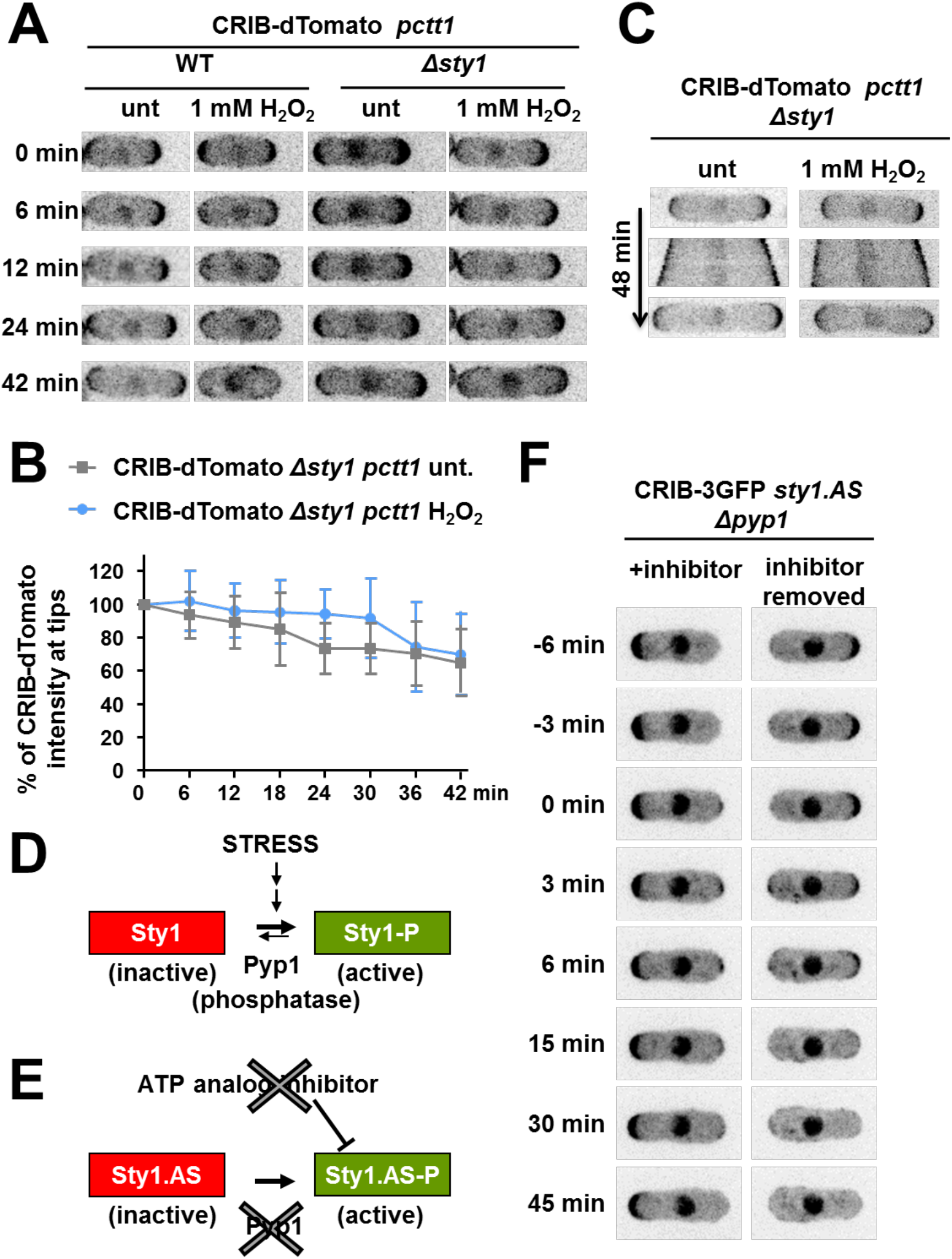
GTP-Cdc42 depolarization from cell tips after oxidative stress depends on Sty1 activation. **(A)** Cdc42-GTP depolarization form cell tips upon H_2_O_2_ imposition is abolished in *Δsty1* cells. Still images showing GTP-bound Cdc42 at several points from time-lapse experiments of YE cultures of strains CS245 (*CRIB-dTomato psty1’::ctt1*) and CS238 (*CRIB-dTomato Δsty1 psty1’::ctt1*) treated or not with 1 mM H_2_O_2_. **(B)** Quantification as in Fig1B of CRIB-tdTomato intensity at cell tips from movies as in A. **(C)** Cell growth is not halted in *Δsty1* cells after stress imposition. Kymographs showing CRIB-dTomato cells from same time-lapse experiments as in A. (**D-E**) Schematic representation of the *sty1*.*AS Δpyp1* mutant. Upon stress conditions, Sty1 activation is counteracted by the phosphatase Pyp1, eventually leading to the shut-off of the cascade. In the *sty1*.*AS Δpyp1* mutant, the absence of Pyp1 leads to constitutively active Sty1 but the presence of an ATP analogue keeps the kinase inactive. In this background, Sty1 can be artificially activated by the removal of the inhibitor. (**F**) Stress-independent activation of Sty1 leads to Cdc42-GTP dispersal from cell tips. Still images showing GTP-bound Cdc42 staining of YE cultures of strain CS147 (*CRIB-3GFP sty1*.*T97A Δpyp1*) growing on the continuous presence of 10 µM of 1NM-PP1 or after inhibitor removal.

In a third strategy, we monitored CRIB-3GFP in cells lacking Pyp1, the main tyrosine phosphatase of active Sty1 (Millar et al., 1995; Shiozaki and Russell, 1995a) (Fig. 3D), and expressing the analog-sensitive mutant Sty1.AS from the endogenous *sty1* gene (Gregan et al., 2007; Zuin et al., 2010). In the Sty1.AS protein, the threonine 97-to-alanine mutation in the kinase ATP pocket facilitates the entry of the bulky inhibitory ATP analog 3MB-PP1, which fully blocks H_2_O_2_-dependent Sty1.AS activity without perturbing wild-type Sty1 (Fig. S3D). Cells lacking Pyp1 display basal activation of Sty1 under basal conditions, but addition of 3MB-PP1 keeps Sty1.AS inactive and downstream Atf1 dephosphorylated, until the analog is withdrawn (Fig. 3E and Fig. S3C). In the presence of the kinase inhibitor, CRIB-3GFP expressed in *Δpyp1 sty1*.*AS* cells localizes to cell tips, but removal of the analog quickly triggers CRIB dispersal from the poles (Fig. 3F), with similar kinetics to the effect of H_2_O_2_ addition to CRIB-3GFP-expressing wild-type cells (compare Fig. S3E and Fig. 1B). From these experiments, we conclude that direct activation of the Sty1 kinase, either genetically or upon stress imposition, is sufficient to redistribute the Cdc42 polarity module from the cell tips to lateral patches.

### Cell length and activity of Cdc42 at the poles inversely correlates to Sty1 activity

The rod-shaped morphology and tip elongation growth of fission yeast has enabled the isolation of mutations advancing or delaying cell cycle progression, based on cell length at division. In fact, a genetic interaction was described more than two decades ago between Sty1 and the cell-cycle machinery (Shiozaki and Russell, 1995a). We have measured cell length and CRIB intensity at cell poles of different MAP kinase pathway mutants before cell division (Fig. 4ABC). While *Δsty1* cells are longer at cell division than wild-type cells, *Δpyp1* (in which the Sty1 pathway is constitutively activated) are shorter (Fig. 4AB). Importantly, CRIB-3GFP expression does not affect cell size in the different backgrounds (Fig. S4A). It is worth noting that *Δatf1* cells are similar to wild-type cells, indicating that the effect of Sty1 in the regulation of cell length is probably independent of transcription (Fig. 4B and Fig. S4A). Regarding active Cdc42 (CRIB-3GFP) at cell tips in the different genetic backgrounds (Fig. 4C), we determined a two-fold increase in CRIB-3GFP fluorescence intensity in *Δsty1* cells; on the contrary, we detected a significant decrease of active Cdc42 at the cell tips of *Δpyp1* cells, enforcing the idea that Sty1 modulates Cdc42 activity at cell tips also at unstressed conditions.

**Figure 4.**
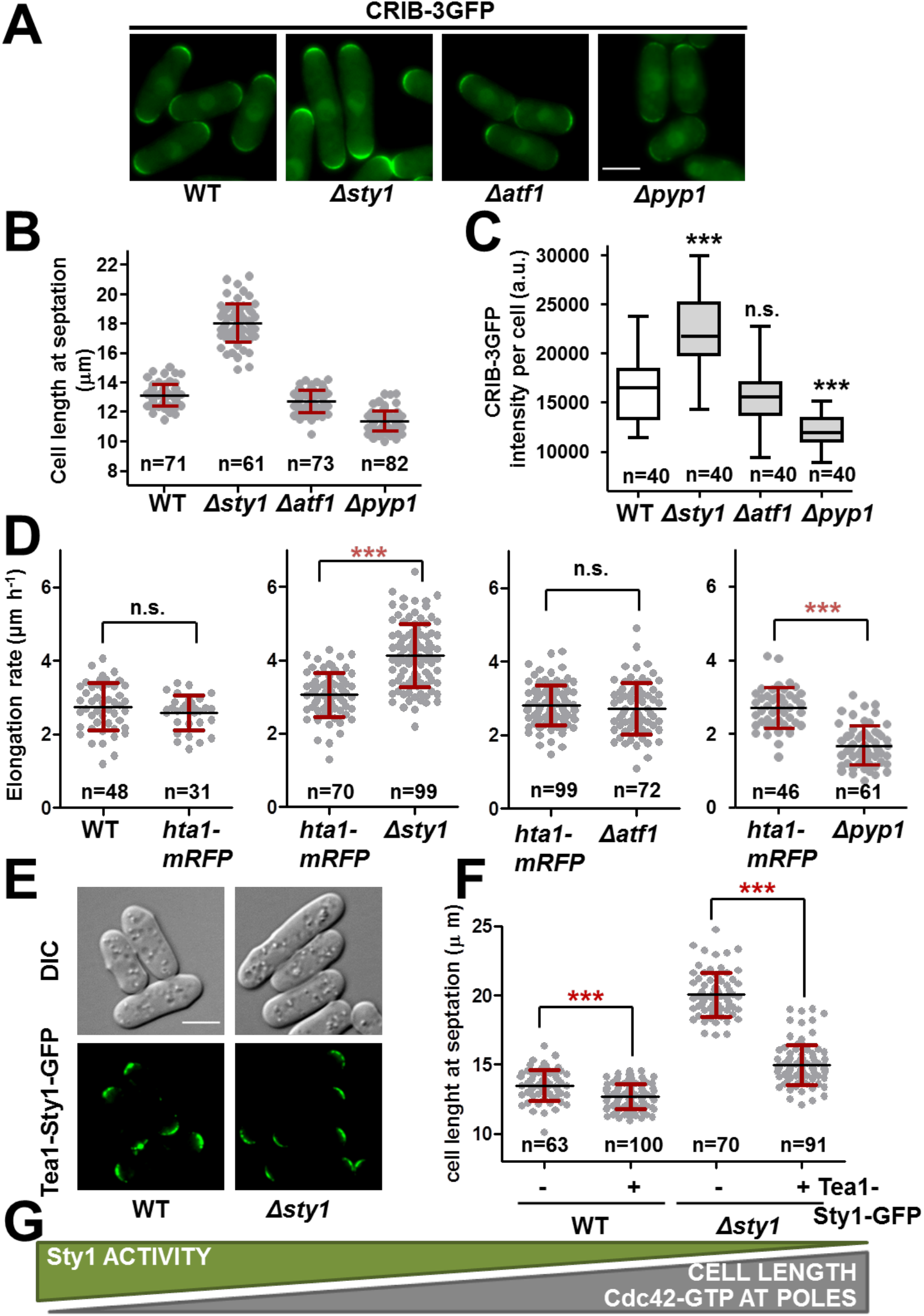
Cell length and activity of Cdc42 at the poles are inversely correlated in mutants of the Sty1 pathway. **(A)** GTP-Cdc42 levels are inversely correlated to Sty1 MAP kinase activation. Representative images of GTP-bound Cdc42 from logharitmic YE cultures of strains PPG65.60 (*CRIB-3GFP*), CS137 (*CRIB-3GFP sty1-1*), CS133 (*CRIB-3GFP Δatf1*) and CS207 (*CRIB-3GFP Δpyp1*). Scale bar: 5 µm. **(B)** Cell length distribution of logarithmic YE cultures of same strains as in A is shown. Mean, SD and number of cells analyzed (n) are indicated. **(C)** Quantification of CRIB-3GFP intensity at tips per cell of the same strains as in A. Whiskers indicating min to max are shown, horizontal line represents mean. The number of cells analyzed (n) is indicated. Statistical significance in regard to wild-type strain was calculated using unpaired t-test (***p-value>0.0001). **(D)** Cdc42 activity is directly related to elongation rate. Elongation rates of MM cultures of 972 (WT), JA1329 (*hta1-mRFP*), AV18 (*Δsty1*), MS98 (*Δatf1*) and EP48 (*Δpyp1*) were assayed. Graphs show mean, SD and number of cells analyzed (n). Statistical significance was determined using unpaired t-test (*** p-value<0.0001). **(E)** Trapping Sty1 to cell tips is sufficient to restore the cell size phenotype. Still images showing the localization of Tea1-Sty1-GFP protein fusion in MM cultures of strains CS309 (*nmt81::tea1-sty1-GFP:leu1+*) and CS310 (*sty1::ura4+ nmt81::tea1-sty1-GFP:leu1+*) are shown. **(F)** Cell length distribution of the same strains as in E is shown. Mean, SD and number of cells analyzed (n) are indicated. Statistical significance was calculated using unpaired t-test (***p-value>0.0001). **(G)** Schematic representation showing the inverse proportionality between Sty1 activity and Cdc42-GTP activity at cell poles/cell length.

A consequence of active Cdc42 deposition at cell tips may be an increase in the growth rate. To facilitate the comparison between different genetic backgrounds, we set up to measure cellular growth rates in co-cultures of wild-type, *Δsty1, Δatf1* and *Δpyp1* strains. To that end, cultured cells were incubated with fluorescein isothiocyanate (FITC)-lectin, which stains cell surface carbohydrates in green, the dye was washed out, and growth proceeded for 90 min to expose unlabeled growth areas, which were counterstained with calcofluor (visualized in red) to highlight new growth surfaces (see Methods for details) (Fig. S4B). In order to compare strains pair-wise, we labeled wild-type cells with a nuclear fluorescent reporter, Hta1-mRFP. This tag does not affect the growth rate of wild-type cells (Fig. 4D, left panel). As observed in Fig. 4D, while wild-type cells elongated at a rate of ∼3 µm/h, *Δsty1* cells did it much faster (≥ 4 µm/h) while *Δpyp1* cells elongated slower (≤ 2 µm/h). As expected, *Δatf1* cells grew at the same rate than wild-type cells.

To completely separate cytosolic from nuclear Sty1 activity, we generated a Sty1 chimera fused to the cell tip-resident Tea1. This Tea1-Sty1-GFP molecule could be readily observed at cell tips in a wild-type or in a *Δsty1* background (Fig. 4E). We also confirmed that this chimera was unable to activate the transcriptional cascade downstream of Sty1 using two different approaches: (i) in the absence of endogenous Sty1, Atf1 was not phosphorylated after treatment of cells with H_2_O_2_ indicating that the chimera lacked Sty1 nuclear functions (Fig. S4C); (ii) *Δsty1* cells expressing Tea1-Sty1-GFP are as sensitive to oxidative stress in liquid growth as cells lacking Sty1 (Fig. S4D). Altogether, these results demonstrate than Sty1 dragged to the cell tips cannot contribute to trigger wild-type tolerance to H_2_O_2_ and lacks nuclear functions. We then tested whether the Tea1-Sty1-GFP chimera, which was constitutively trapped at cell tips, contributes to regulate cell length at septation. As can be observed in Fig. 4F, expressing Sty1 at cell tips slightly, but significantly, reduced cell length at division on wild-type cells. More noticeably is the effect in a *Δsty1* background, where having Sty1 tethered at cell tips reduced the length at septation from ≥20 µm to ∼15 µm (Fig. 4FG).

### Inhibition of GEF activity mediates the stress-dependent effect on growth polarity

In search for downstream effectors of Sty1 in its inhibitory role on polarity, we considered that Sty1 inactivation or delocalization of the two S. pombe GEFs, Gef1 or Scd1, could mediate the stress-dependent inhibition of Cdc42-GTP. We tested whether the exit from cells tips of these GEFs upon stress could cause Cdc42 inactivation. Cells lacking Scd1 are rounded due to its essential role in establishing sites of polarity, and display weak patches of active Cdc42 (Fig. S5A). In an attempt to recover cell polarity of *Δscd1* cells through the artificial tethering of either Gef1 or Scd1 to cell tips, we expressed in this background chimeras of Gef1 or Scd1 fused to Tea1. As shown in Fig. 5A, both GEF-based chimeras, Tea1-Scd1-GFP and Tea-Gef1-GFP, when expressed in cells lacking Scd1 are localized to cell tips and are able to promote the accumulation of CRIB-3mCherry at the growing poles. They fully suppress the rounded phenotype of strain *Δscd1*, both in length (Fig. 5B) and in width (Fig. S5B). With this constitutively tethered GEF-based chimeras in cells lacking Scd1, we tested the effect of stress on polarized growth. Probably due to the different conformation of the large chimeras, the fluorescence levels of Tea1-Scd1-GFP and Tea1-Gef1-GFP are different, but both proteins are retained at cell poles after stress imposition, as expected (Fig. 5CD). H_2_O_2_ causes dispersal of active Cdc42 from the cell tips to lateral patches in *Δscd1* cells expressing Tea1-Scd1-GFP (Fig. 5CE), indicating that the exit of Scd1 from cell poles (see Fig. 2A) is not the cause of reduced Cdc42 activity upon stress. Strikingly, *Δscd1* cells expressing Tea1-Gef1-GFP are blind to stress imposition: CRIB-3mCherry remains at cell tips and lateral patches are not formed (Fig. 5CD). Importantly, inversion of the fluorescent tags (Tea1-Gef1-mCherry and CRIB-3GFP) in *Δscd1* cells (Fig. S5C) or in a *Δgef1* background (Fig. S5D) also yields stress-insensitive strains. This experiment demonstrates that cells displaying a rod-like phenotype thanks to the constitutively tip-anchored Tea1-Gef1 are not sensitive to stress inhibition, and suggests that removal of Gef1 from cell tips could be one of the molecular events leading to the stress-dependent Cdc42 inactivation in wild-type cells. However, since cells lacking Gef1 do not block Cdc42 depolarization we speculate that other mechanism must collaborate in the stress-dependent Cdc42 inhibition (Fig. 2B).

**Figure 5.**
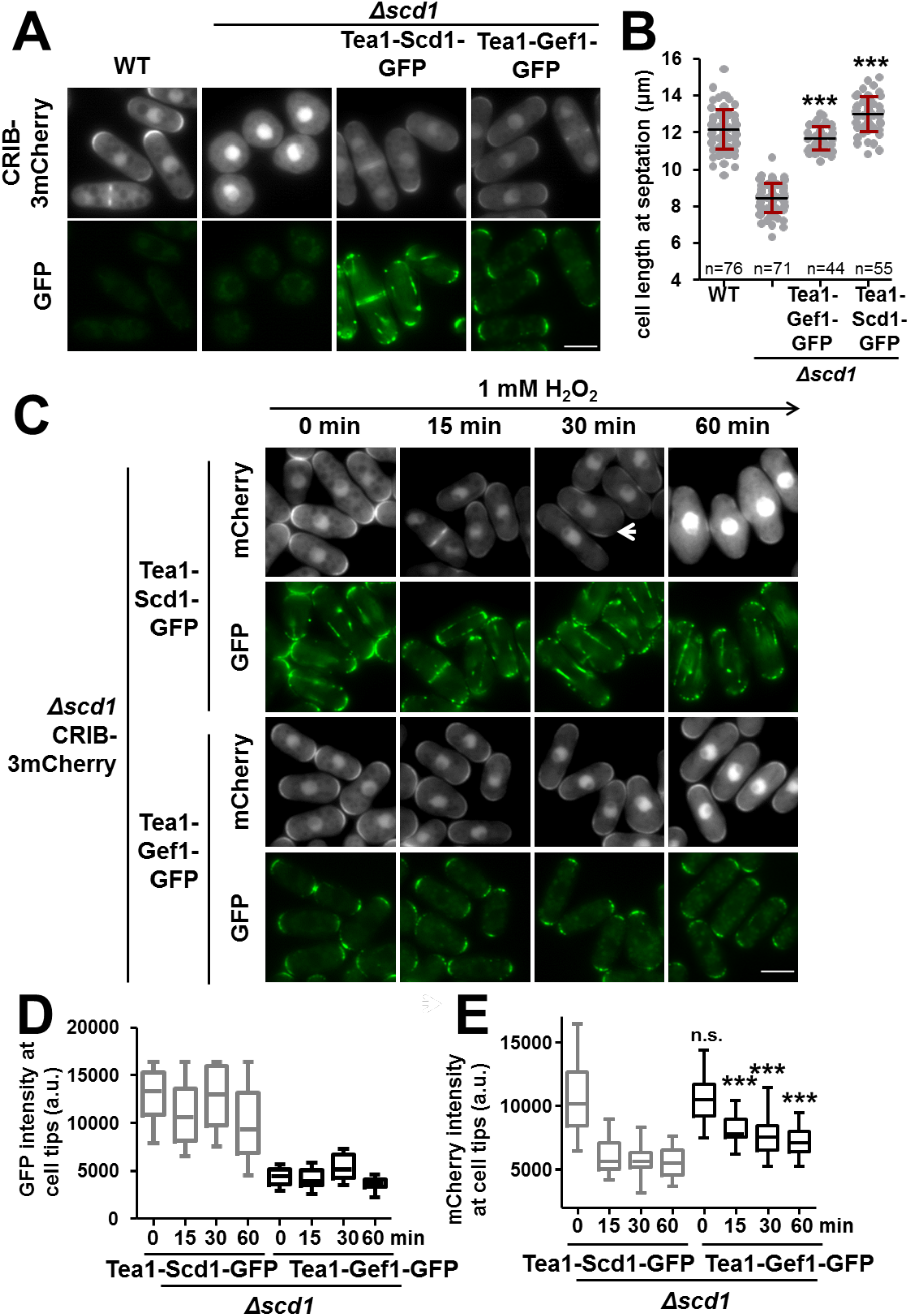
Inhibition of GEF activity mediates the stress-dependent effect on polarity. **(A)** Trapping GEFs to cell tips restores cell length and GTP-Cdc42 levels in *Δscd1* cells. Representative images showing CRIB-3mCherry and GFP staining of MM cultures of CS357 (*CRIB-3mCherry*), CS359 (*CRIB-3mCherry Δscd1*), CS365 (*CRIB-3mCherry Δscd1 nmt81::tea1-scd1-GFP*) and CS360 (*CRIB-3mCherry Δscd1 nmt81::tea1-gef1-GFP*). Scale bar: 5 µm. **(B)** Cell length distribution at septation of the same strains as in A. Mean, SD and number of cells analyzed (n) are indicated. Statistical significance was calculated using unpaired t-test in regard to *CRIB-3mCherry Δscd1* mutant (***p-value>0.0001). **(C)** Cells expressing Tea1-Gef1, but not Tea1-Scd1, retain GTP-Cdc42 activity after stress imposition. Representative images showing CRIB-3mCherry and GFP staining of MM cultures of CS360 (*CRIB-3mCherry Δscd1 nmt81::tea1-gef1-GFP*) and CS365 (*CRIB-3mCherry Δscd1 nmt81::tea1-scd1-GFP*) treated with 1 mM H_2_O_2_ for the indicated times. GFP fluorescent levels are not equalized between the two strains. Scale bar: 5 µm. **(D)** Quantification of Tea-GEF-GFP intensity at cell tips of the same strains as in C. Whiskers representing min to max are shown, horizontal line indicates mean. Statistical significance of CS365 versus CS360 was calculated using unpaired t-test (n.s. non-significant, *p-value>0.05, ***p-value>0.0001). At least forty tips were analyzed for each condition within at least two independent biological replicates. **(E)** Quantification of CRIB-3mCherry intensity at cell tips of the same strains as in C. Represented as in D.

### Cells devoid of the Cdc42 GAPs Rga3 and Rga6 are insensitive to stress-dependent cell polarity inhibition

In the search for additional downstream effectors of Sty1 in the polarity module, we then interrogated the putative GAPs of Cdc42, since these activators of the GTP-to-GDP hydrolysis are obvious switch-off molecules of the GTPase. As shown by several laboratories, Rga4 and Rga6 localize to cell sides, and contribute to the spatial restriction of active Cdc42 at cell poles (Revilla-Guarinos et al., 2016; Tatebe et al., 2008), while Rga3 has been recently described to co-exist with GTP-Cdc42 at growth sites (Gallo Castro and Martin, 2018). We first monitored the localization of Rga3, Rga4 and Rga6 after environmental stress. To do so, we co-expressed CRIB-3mCherry with GFP-tagged versions of each GAP protein. Rga4-GFP remains in lateral patches before and after stress imposition (Fig. 6A), and *Δrga4* cells are sensitive to apical growth inhibition after H_2_O_2_ stress (Fig. S6A), discarding any role of this GAP in Sty1-dependent Cdc42 inhibition. Rga6-GFP still localizes at the lateral membranes after H_2_O_2_ imposition, but some clusters also diffuse into the cell tips, as reported before upon LatA treatment (Fig. 6A) (Revilla-Guarinos et al., 2016). Moreover, in *Δsty1* cells Rga6-GFP is excluded from cell tips even after LatA treatment (Fig. 6B), indicating that Rga6 localization may be directly regulated by Sty1 kinase. Finally, Rga3-GFP remains at cell tips after treatment with H_2_O_2_ (Fig. S6B), as reported before with LatA (Gallo Castro and Martin, 2018). Therefore, two GAPs are present at cell tips after stress (Rga3 and Rga6) and therefore are candidates to trigger Cdc42 inactivation (Fig. 6C).

**Figure 6.**
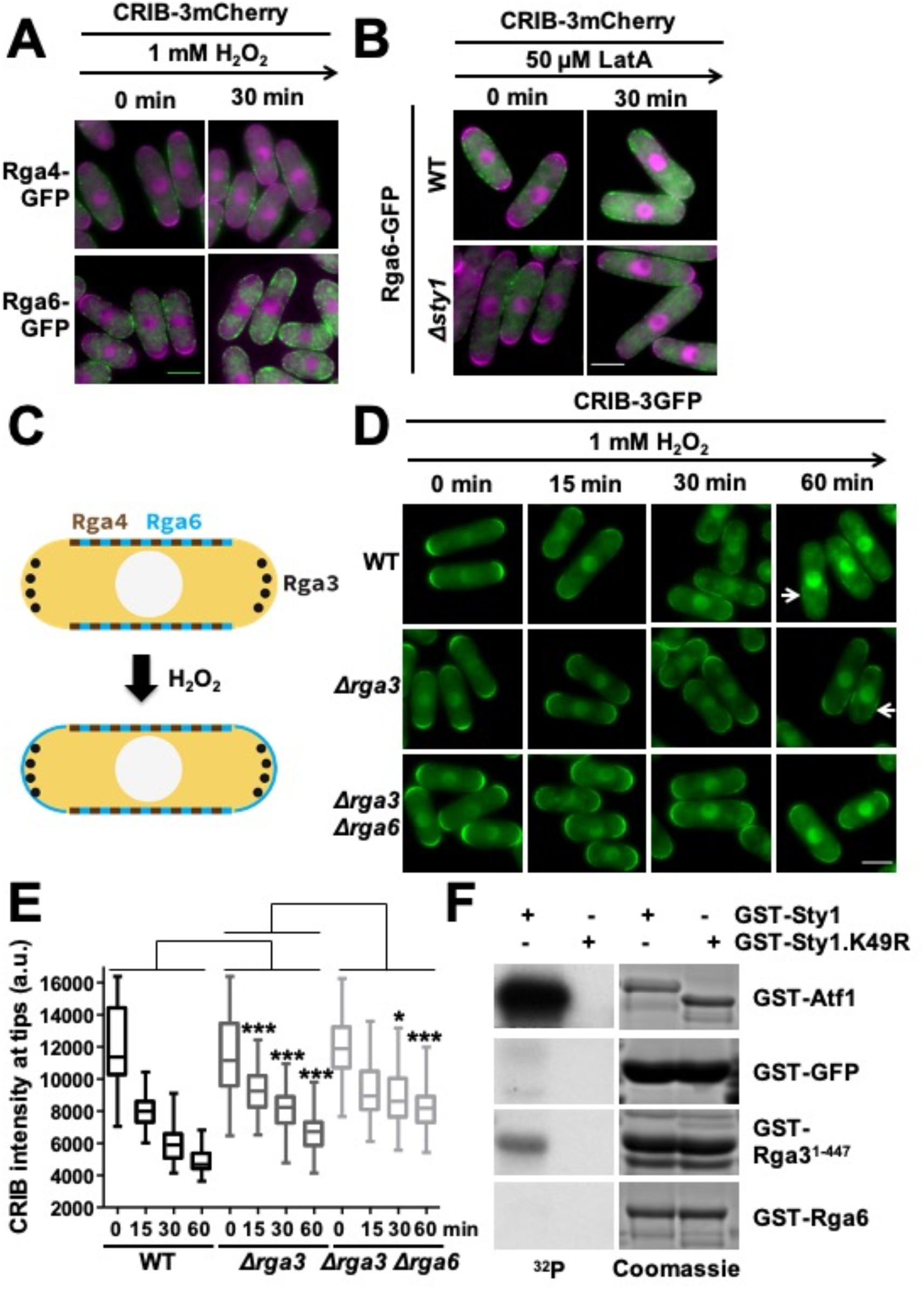
Rga3 is a direct target of Sty1 kinase. **(A)** Rga6 spreads to cell tips after stress imposition. Representative images showing merged staining of GTP-bound Cdc42 (in magenta) and GFP-tagged versions of Rga4 and Rga6 (in green). YE cultures of CS361 (*CRIB-3mCherry rga4-GFP*) and CS362 (*CRIB-3mCherry rga6-GFP*) were treated with 1 mM H_2_O_2_ for the indicated time. Scale bar: 5 µm. **(B)** Rga6 does not decorate cell tips after stress imposition in cells lacking Sty1. Representative images showing merge staining of GTP-bound Cdc42 (in magenta) and GFP-tagged version of Rga6 (in green). YE cultures of CS362 (*CRIB-3mCherry rga6-GFP*) and CS369 (*CRIB-3mCherry rga6-GFP Δsty1*) were treated with 50 μM LatA for 30 min. Scale bar: 5 µm. **(C)** Schematic representation of GAPs cellular localization before and after H_2_O_2_ imposition. **(D)** GTP-Cdc42 depolarization is noticeably reduced in cells lacking Rga3 and Rga6. Representative images of GTP-bound Cdc42 from YE cultures of PPG65.60 (*CRIB-3GFP*), CS244 (*CRIB-3GFP Δrga3*) and CS315 (*CRIB-3GFP Δrga3 Δrga6*) treated with 1 mM H_2_O_2_ for the indicated times. White arrows indicate the presence of lateral patches. Scale bar: 5 µm. **(E)** Quantification of CRIB-3GFP intensity at cell tips of the same strains as in D. Representation as described in 5D. **(F)** Rga3 is directly phosphorylated by Sty1 *in vitro*. The indicated substrates were assayed against GST-Sty1 or the kinetic dead version GST-Sty1.K49R using γ-^32^P-ATP.

We then tested the effect of single and double deletion of GAP-coding genes on stress-dependent inactivation of cell polarity. While *Δrga6* cell expressing CRIB-3GFP display H_2_O_2_ inactivation kinetics similar to wild-type cells (Fig. S6CD), cells lacking Rga3 retain active GTP-Cdc42 at cell tips 60 min after stress imposition (Fig. 6D). In this strain background, the main GEF Scd1 is still dispersed from the tips upon stress (Fig. S6E). Strikingly, the growth poles of *Δrga3 Δrga6* cells retain higher Cdc42 activity than single *Δrga3* cells and lateral patches are not present after H_2_O_2_ imposition (Fig. 6DE). We conclude that cells lacking the Cdc42 GAPs Rga3 and Rga6 lack the hallmarks linked to stress-dependent cell polarity inhibition (dispersal of CRIB-3GFP from poles and appearance of GTP-Cdc42 lateral patches).

We tested whether some of these targets are direct substrates of Sty1 using recombinant proteins in *in vitro* kinase assays. First of all, we mapped the canonical S/TP sites in Rga3 and Rga6 proteins. As shown in Fig. S6F, most of the sites in Rga3 are concentrated in the N-terminus. On the other hand, Rga6 sites are dispersed along the protein sequence. We purified a GST-tagged fragment of Rga3 and the full length version of Rga6, and we tested their phosphorylation by wild-type GST-Sty1 or its catalytically-dead version GST-Sty1.K49R. As positive and negative controls we used GST-Atf1 and GST-GFP, respectively. As shown in Fig. 6F, a GST-tagged fragment of Rga3 carrying most putative MAP kinase sites, was clearly phosphorylated by Sty1 but not by the catalytically-dead version GST-Sty1.K49R. We propose that Rga3 is a direct target of Sty1, and that this phosphorylation activates the GAP to initiate Cdc42 depolarization at cell tips after stress imposition. In summary, our data suggest that the activation of Rga3 GAP, in combination with Gef1 depolarization promotes the transient inhibition of cell growth and polarity upon stress.

## DISCUSSION

Adaptation to stress includes multiple cellular responses, most of them transcriptional or resulting in cell cycle inhibition. Signalling cascades often mediate a variety of effects through modification of diverse downstream effectors, such as transcription factors or cell cycle regulators. Here, we describe the effect of the MAP kinase Sty1 on cell polarity. By directly regulating factors of the polarity module at the cell tips, the H_2_O_2_-activated kinase alters the equilibrium between GAPs and GEFs and transiently inactivates Cdc42, the driving force of the polarity module, till growth conditions are favourable again. When stress is applied, three related phenomena are observed: active GTP-Cdc42 is dispersed from cell tips, lateral patches of GTP-Cdc42 are formed and cellular growth rates are fully inhibited.

Lateral patches after GTP-Cdc42 dispersal from cell tips are observed in a variety of Sty1-activating conditions, including LatA treatment (Bendezu and Martin, 2011; Hercyk et al., 2019; Mutavchiev et al., 2016), nutrient depletion (Chen et al., 2019), heat shock (Vjestica et al., 2013), osmotic stress (Haupt et al., 2018) and oxidative stress, as reported here. However, which actors play a role on the formation of these GTP-Cdc42 lateral patches remains unsolved. A recent report has shown the existence of a positive feedback loop in fission yeast causing the deposition and enrichment of GTP-Cdc42 sites at cell tips based on the quaternary complex Cdc42-Pak1-Scd2-Scd1 (Lamas et al., 2020), similar to that extensively studied in budding yeast. In addition, the group of Martin has shown that Scd1 can be artificially substituted by Gef1 in the quaternary complex, indicating that GEF activity is the key contribution of the positive feedback (Lamas et al., 2020). Our data shows that the quaternary complex is broken at cell tips upon stress imposition and the scaffold protein Scd2, the secondary GEF Gef1 and the downstream effector Pak1 are present in GTP-Cdc42 lateral patches. Interestingly, we could not detect the presence of the main GEF Scd1 and the activation of Cdc42 at these sites relies uniquely on Gef1, as also reported in response to LatA (Hercyk et al., 2019). We also show that Scd2 is dispensable for Cdc42 activation at the lateral patches, suggesting that the scaffolding activity is not required for Cdc42-Gef1 interaction. Even though Cdc42 effector proteins, including the formin For3 and components of the exocyst, are located in the lateral patches (Bendezu and Martin, 2011), Cdc42 activation at these sites does not lead to any observable morphological changes. Probably, co-localization with GAP proteins and/or the presence of downstream inhibitors limit further downstream activation of the Cdc42 cascade.

Negative feedback loops limit the number and/or the size of the Cdc42-driven polarity sites under physiological conditions. The concentration of polarity factors at the tips can increase and decrease in an oscillatory behaviour caused by positive and then negative regulators of the circuit, respectively (Das et al., 2012). For instance, phosphorylation of the budding yeast GEF, CDC24, by the downstream effector kinase PAK1 is required to limit the activity of the module (Howell et al., 2012; Kuo et al., 2014). Here, we propose that Sty1 pathway is a key orchestrator of a negative regulatory event on the Cdc42 polarity module, and by doing so it links the physiological behaviour of the module with environmental signals. Our experiments suggest that Sty1 limits the amount of active Cdc42 at cell tips by activating GAPs and inhibiting GEFs, to promote cell growth arrest.

Upon stress imposition, a rising on the GTPase activity could unbalance the GTP/GDP equilibrium of Cdc42 towards the inactive state. Indeed, cells lacking Rga3 display a slower Cdc42 depolarization from cell tips after H_2_O_2_ imposition. Importantly, basal GTP-Cdc42 concentration at cell tips is not affected in *Δrga3* cells, indicating that slower depolarization is not due to a higher initial concentration of GTP-Cdc42. Our in vitro kinase assays showed that Rga3 is directly phosphorylated by Sty1, indicating that Rga3 is one of the targets linking the MAP kinase pathway to the Cdc42 polarity module. Unlike the other Cdc42 GAPs, Rga3 co-localizes with active Cdc42 at cell tips before and after stress (Gallo Castro and Martin, 2018). Interestingly, budding yeast Cdc42 GAPs are also regulated by phosphorylation by cyclin-dependent kinases inhibiting GAP activity (Knaus et al., 2007; Sopko et al., 2007). Our experiments suggest that Rga3 functions as a “stress” GAP: during unperturbed conditions it remains inactive and co-localizes with GTP-Cdc42; upon phosphorylation by Sty1 it rapidly promotes Cdc42 GTPase activity. Inactivation of Cdc42 by Rga3 requires the synergistic participation of Rga6, since *Δrga3* cells still display some depolarization as shown by the formation of lateral patches. Our data demonstrate that Rga6 moves towards cell tips after oxidative stress and *Δrga3 Δrga6* cells do retain higher concentration of GTP-Cdc42 at cell tips. We propose that Rga6 collaborates with Rga3 in the Cdc42 switch-off at later stages of the stress response.

Since CRIB signal at cell tips is still reduced in the double GAP mutant, we reasoned whether GEF proteins could be inhibited after stress imposition. Some evidence indicates that Scd1, which acts as the main GEF during interphase, is no longer active after stress imposition: (i) active Cdc42 on *Δrga3* cells is entirely dependent on Gef1; (ii) trapping Scd1 to cell tips does not affect Cdc42 depolarization, but trapping Gef1 does. However, which are the signalling pathways regulating GEFs inhibition remains unclear. On the other hand, Gef1 remains active upon stress imposition and its artificial anchoring at the cell poles prevents Cdc42 depolarization. This data indicates that stress-dependent Scd1 inhibition and Gef1 delocalization from the cell tips further impact the activity of Cdc42 polarity module upon oxidative stress. However, we can not determine whether these events are directly regulated by Sty1.

In conclusion, our experiments with different genetic backgrounds suggest that stress-activated Sty1 alters the physiological GAP/GEF equilibrium on Cdc42 activity, causing a transient inactivation of cell growth.

We have shown here that the elongated phenotype of cells lacking Sty1 is, at least partly, due to a hyperactivation of the Cdc42 polarity module. Using mutants with different kinase activity levels, we have shown that Sty1 activity is inversely proportional to GTP-Cdc42 levels at cell tips. Consequently, it is also inversely correlated with cell elongation rates and, eventually, cell length at septation. Anchoring Sty1 to cell tips rescues the cell length phenotype, suggesting that targets of Cdc42 regulation at unstressed conditions are also tip-localized. We favor the view that the stress and cell size regulation of Cdc42 by Sty1 is undergone through the same target protein(s) and that Cdc42 activity may be controlled by Sty1-mediated regulatory mechanisms during unperturbed mitotic growth: to initiate growth at the new pole, to proceed with cytokinesis or for mating during sexual reproduction. Indeed, cells lacking Sty1 have severe defects associated with those processes, which could be or not connected to Cdc42 activity. Further work will be required to fully understand the relationship between Sty1 and unperturbed cell cycle events, and to extrapolate our main finding, the connection of the stress-dependent pathway with cell growth and polarity, to other eukaryotic model systems.

## EXPERIMENTAL PROCEDURES

### Growth conditions, genetic manipulations, strains and reagents

Cells were grown in rich medium (YE) or minimal medium (MM) at 30°C as described previously (Alfa et al., 1993). Strains under the control of thiamine-repressible *nmt81* promoter were grown under the presence of 0.02 mg/ml thiamine until saturation. In order to induce gene expression, a small volume of this pre-inoculum was diluted in MM media and cells were let to grow for 16 hours. Origins and genotypes of strains used in this study are outlined in Supplementary Table 1, and most of them were constructed by standard genetic methods. *Escherichia coli* DH5α was used as a host for propagation and construction of the plasmids used in this study. *E*.*coli* cells were grown in LB medium supplemented with 0.1 mg/ml ampicillin. Plasmid p422’ (Paulo et al., 2014), containing the *ctt1* gene under the control of the constitutive *sty1* promoter, was linearized and integrated by homologous recombination at the *leu1-32* locus of different strain backgrounds. To create Tea1 protein fusions, we PCR-amplified genomic *tea1* and cloned it into an episomal *nmt81*-driven background, yielding p696.81. Next, a PCR product containing a linker-GFP coding sequence was amplified from pARC2080, which was kindly provided by Paul Nurse (Kelly and Nurse, 2011) and cloned into the previous plasmid generating the episomal plasmid p697.81. The resulting *nmt81-tea1-(linker)-GFP* was cloned into an integrative background yielding plasmid p701.81’. A BamHI restriction site located in frame between Tea1 and the (linker)-GFP coding sequence (CDS) was used to clone PCR-amplified *sty1* CDS, yielding p712.81’, and *gef1* CDS, yielding p717.81’. As *scd1* CDS contains an internal BamHI site, compatible ends were generated by BglII digestion and cloned into BamHI-linearized p701.81’, yielding the plasmid p747.81’. All these constructions were linearized and integrated by homologous recombination at the *leu1-32* locus of different strain backgrounds. Constructions were checked by sequencing. *tea1* CDS contains I359V and T983A mutations that were generated during PCR amplification. Tea1-mCherry constructions were created using a similar approach. *tea1* CDS was freed from p696.81 and cloned into an integrative vector, yielding p734.81’. mCherry CDS was amplified by PCR and cloned into the previous plasmid using BamHI/SmaI, yielding p748.81’. The resulting plasmid codes for integrative *nmt81*-driven Tea1-mCherry. As previously described, a BamHI restriction site located in frame between *tea1* and *mCherry* was used to clone *gef1* CDS, yielding p749.81’. To generate the GST-tagged protein versions used on the kinase assay *sty1* wild-type and *sty1*.*K49R* ORF were amplified by PCR and cloned into pGEX-4T-1 using BamHI/SmaI yielding p109 and p109.K49R. *wis1*.*DD* (S469D, T473D) was amplified by PCR and cloned into pGEX-4T-1 using XhoI/SmaI, yielding p773. *rga6* ORF was PCR-amplified and cloned into pGEX-4T-1 using EcoRI/SmaI yielding p759. *atf1* full-lenght and the fragment from aminoacid 1 to 447 of *rga3* ORF were amplified by PCR and cloned into pGEX-4T-1 using BamHI/SmaI and yielding p771 and p755, respectively. For the negative control, the linker+GFP sequence described above was freed using BamHI/SmaI and cloned into pGEX-4T1 yielding p772. pAV0756 coding for stable-integrative version of CRIB-3mCherry was obtained from the Japanese National BioResource Project (NBTP; ttp://yeast.nig.ac.jp/yeast) and cloned at the wild-type *his5* locus as described elsewhere (Vjestica et al., 2020). H_2_O_2_ and LatA was purchased from Sigma (H1009 and L5163, respectively), the ATP-analogues 1-NM-PP1 and and 3-MB-PP1 were obtained from Toronto Research Chemicals (A603003; A602960).

### Microscopy techniques and image processing

All microscopy experiments were done with cell cultures at logarithmic phase, at an OD600 of ∼0.5, in either YE or MM, as indicated in each figure legend. For time-lapse experiments, 8-chamber coverglass slides (155411, ThermoScientific) were coated with 1 mg/ml of lectin from glycine max soybean (L1395, Sigma) and left for 30 min. Excess of lectin was washed with the appropriate medium and 200 µl of cell culture was added to each chamber and left to settle for 2 min. Next, cells were washed once with the appropriate media and the same volume of pre-warmed fresh media was added. The preparations were placed at the microscope chamber at 30°C, except for experiments involving pretreatment with 3MB-PP1. In that case, media containing the drug was added and the chamber was left for 60 min in a 30°C incubator. Time-lapse experiments were performed using spinning disk confocal microscope (Revolution XD Andor Technology) with a Plan Apochromat 100x, 1.45 NA objective equipped with a dual-mode electron-modifying charge-coupled device camera (iXon 897 E; Andor Technology). Seventeen z-stacks along 4 µm were acquired every 3 min for CRIB-GFP cells and every 6 min for CRIB-tdTomato cells. iQ Live Cell Imaging software (Andor Technology) was used for image acquisition. For the addition of either LatA or H_2_O_2_ during imaging, media was removed carefully using a 1 ml transfer pipette (13469118, FisherBrand) and the same amount of media (200 µl) containing the appropriate compound was added. This process was completed within a 3-min interval so the acquisition timing was not disrupted. Fiji (Image J, National Institutes of Health) (Schindelin et al., 2012) was used for image processing. Still images of time-lapse experiments are presented as z-stacks at maximum intensity. In order to quantify Cdc42 depolarization from time-lapse experiments (as in Fig. 1B, 3B, S3A, S3D and S3F), the mean value of each cell cytoplasm was subtracted from the image in order to obtain pure CRIB signal at tips. Tip signal was narrowly selected using the square tool and integrated density of each tip was measured along the different time points. To obtain CRIB signal per cell, the values of both tips were added and the percentage of CRIB intensity was calculated by referencing all the values to time zero. Kymographs were manually generated using Fiji imaging software. Briefly, the stack of interest was aligned using StackReg plugin. Then a montage consisting of different time-points was done from a horizontal rectangle of 5 μm located along the cell cortex. For DIC and conventional fluorescence microscopy images, we used a Nikon Eclipse 90i microscope equipped with differential interference contrast optics, a PLAN APO VC 100× 1.4 oil immersion objective, an ORCA-II-ERG camera (Hamamatsu) image acquisition software Metamorph 7.8.13 (Gataca Systems) and a LED illumination Cool LED pE-300lite. For quantification of CRIB intensity (Fig. 4C, 5E, S5D and S5G) at cell tips in this case a different approach was used: we draw a straight line using Fiji centered along the “x” axis of the cell (from one cell tip to the other), then a histogram was generated. The peak values of fluorescence intensity in each tip were added obtaining CRIB intensity per cell. For cell length and cell width measurements cells were stained with 0.1 mg/ml calcofluor (Sigma, F3543) and examined under the microscope. Only cells with visible septum were analyzed. All cell measurements were performed manually using the line tool of Fiji software. Graphs and statistical analysis were performed with Prism (GraphPad Software).

### Growth zones staining using FITC

Novel growth zones were measured as described previously (Revilla-Guarinos et al., 2016).

Briefly, cells were grown to an OD_600nm_ of ∼0.3 and two different strains were mixed at equal parts in MM media containing 5 μg/ml of FITC-lectin (L9381, Sigma-Aldrich). After 15 min of incubation, cells were pelleted by mild centrifugation and washed twice with MM. Then cells were further incubated with stain-free pre-warmed media for 90 min. Finally, cells were counterstained with 0.1 mg/ml calcofluor white, observed under the microscope and measured as described.

### H_2_O_2_ sensitivity assay

In order to monitor sensitivity to stress imposition, growth curves were performed as described elsewhere (Calvo et al., 2009). Briefly, MM cultures at an initial OD_600nm_ of 0.1 were treated or not with LatA or H_2_O_2_ and inoculated by duplicate in 96-well non-coated polystyrene microplates with an adhesive plate seal. Plates were incubated in a Power Wave microplate scanning spectrophotometer (Bio-Tek) at 30°C with continuous shaking. The OD_600nm_ was automatically recorded every 10 min for the indicated hours using Gen5 software.

### TCA extracts and immuno blot analysis

Modified trichloroacetic acid (TCA) extracts as previously described (Jara et al.; 2007), were separated by SDS-PAGE and detected by immunoblotting with polyclonal anti-Atf1 (Sanso et al., 2008), polyclonal anti-Sty1 (Jara et al.; 2007), monoclonal anti-GFP (Takara) and monoclonal anti-tubulin (T5168, Sigma-Aldrich) antibodies.

### Protein production in E.coli and *in vitro* kinase assay

In order to produce and purify GST-tagged proteins, the protease deficient *E. coli* strain FB810 was transformed with the corresponding plasmids. A liquid culture of 200 ml was grown up to OD_595nm_ 0.6, then 100 μM of IPTG was added to the culture and further incubated during 20 h at 18°C. Cells were collected and resuspended in STET buffer [50 mM Tris HCl pH 8.0, 1 mM EDTA, 150 mM NaCl, 5% Triton X-100]. Cell lysis was obtained by 5 pulses of sonication of 15 sec at 40% amplitude. Soluble fraction was prepared by centrifugation at 10,000 rpm for 10 min at 4°C. A volume of 2.5 ml of soluble fraction was incubated with 100 μl of Glutathione Sepharose beads (GE Healthcare, 17-0756-01) for 1 h at 4°C. Beads were washed three times with NET-N buffer [20 mM Tris-Hcl pH 8.0, 1 mM EDTA, 100 mM NaCl, 0.5% NP-40]. A final washing in Elution Buffer [100 mM Tris-HCl pH 8.0, 120 mM NaCl] was performed. Finally the desired protein was eluted form the beads by incubation with 100 μl of Elution Buffer containing 6 mg/ml of GSH (Sigma, 4251) during 30 min at 4°C. Recombinant proteins were checked and quantified in a polyacrylamide gel stained with Comassie Brilliant Blue (Amresco, 0472). The proteins were stored at −80°C. In order to perform the kinase assay two sequential enzymatic reactions were prepared. First of all, GST-Sty1 kinase should be activated by pre-incubation with its MAP2K GST-Wis1.DD. To prepare reaction A, 1 μg of either GST-Sty1 or GST-Sty1.K49R was mixed with 0.3 μg of GST-Wis1.DD in 5x Kinase buffer [250 mM Tris-HCl pH 7.5, 250 mM KCl, 50 mM MgCl_2_, 2 mM DTT] containing 0.5 mM of freshly added rATP (Promega, E6011) in 10-μl reactions. Reaction A was incubated at 30°C for 5 min. Then it was mixed with reaction B which contain 5 μg of the GST-substrate in 5x Kinase buffer containing 2 μCi γ-^32^P-ATP (PerkinElmer, BLU002Z250UC) in 10-μl reactions. The mix was further incubated at 30°C for 20 min. The enzymatic reaction was stopped by adding SDS-PAGE sample. Proteins were boiled and separated into a polyacrylamide gel. Gels were dried overnight and the signal of γ-^32^P-ATP was detected by film exposition.

### Sequence analysis

Protein domains as in Fig S6G were obtained using a simple modular architecture research tool (SMART) (http://SMART.embl-heidelberg.de).

Canonical S/T-P phosphorylation sites were mapped manually.

## ACKNOWLEDGEMENTS

We thank Paul Nurse and Fulvia Verde for providing strains and plasmids. We are especially thankful to Aleksandar Vjestica and Sophie Martin for the generation and generous sharing of fission yeast genetic tools. We thank Margarita Cabrera and Javier Encinar del Dedo for technical support with image analysis. This work is supported by the Ministerio de Ciencia, Innovación y Universidades (Spain), PLAN E and FEDER (PGC2018-093920-B-100 to E.H. and PGC2018-097248-B-100 to J.A.). The Oxidative Stress and Cell Cycle group is also supported by Generalitat de Catalunya (Spain) (2017-SGR-539) and by Unidad de Excelencia María de Maeztu, funded by the AEI (CEX2018-000792-M) (Spain). C.S.-C. is recipient of a María de Maeztu predoctoral fellowship from the Ministerio de Economía y Competitividad (Spain). E.H. is recipient of an ICREA Academia Award (Generalitat de Catalunya, Spain).

## AUTHOR CONTRIBUTIONS

C.S.-C. performed most experiments. M.C was involved with GST-proteins purification. R.G.-M. performed some microscopy experiments. C.S.-C., J.A., P.P. and E.H. analyzed the data. E.H. and C.S.-C. wrote the manuscript.

