## Supplemental Information for "Stress-dependent inhibition of cell polarity through unbalancing the GEF/GAP regulation of Cdc42"

| Strain | Genotype | Origin |
|---|---|---|
| 972 | $h^-$ | (Leupold, 1970) |
| AV18 | $h^-$ sty1::kanMX6 | (Zuin et al., 2005) |
| CS133 | $h^?$ ppak1'::CRIB-3GFP::ura4+ atf1::natMX6 | This study |
| CS136 | $h^-$ ppak1'::CRIB-3GFP::ura4+ sty1.T97A | This study |
| CS137 | $h^?$ ppak1'::CRIB-3GFP::ura4+ ura4D-18 sty1-1 leu1-32 | This study |
| CS147 | h? ppak1'::CRIB-3GFP::ura4 ura4-D18 pyp1::natMX6 sty1.T97A | This study |
| CS207 | $h^+$ pyp1::kanMX6 ppak1'::CRIB-3GFP::ura4+ ura4-D18? leu1-32 | This study |
| CS208 | $h^?$ pact1::.lifeact-mCherry::leu1+ pak1::CRIB-3GFP::ura4+ leu1-32 ura4-D18 | This study |
| CS235 | $h^-$ rga3-GFP | This study |
| CS238 | $h^+$ ppak1::CRIB-linker-dTomato::ura4+ sty1::natMX6 psty1'::ctt1::leu1+ ura4-D18 leu1-32 ade6-M210 | This study |
| CS244 | $h_?$ ppak1::CRIB-3GFP::ura4+ rga3::kanMX6 ura4-D18 leu1-32 | This study |
| CS245 | $h^+$ ppak1::CRIB-linker-dTomato::ura4+ psty1'::ctt1::leu1+ ura4-D18 leu1-32 ade6-M210 | This study |
| CS281 | $h^+$ ppak1::CRIB-3GFP::ura4+ scd2::kanMX6 ura4-D18 leu1-32 | This study |
| CS309 | $h^-$ nmt81'::tea1-sty1-GFP::leu1+ leu1-32 | This study |
| CS310 | $h^-$ nmt81'::tea1-sty1-GFP::leu1+ sty1::ura4+ ura4-D18 leu1-32 | This study |
| CS315 | $h^+$ ppak1'::CRIB-3GFP::ura4+ rga3::natMX6 rga6::kanMX6 ura4-D18 leu1-32 | This study |
| CS316 | $h^?$ ppak1'::CRIB-3GFP::ura4+ rga3::natMX6 gef1::kanMX6 ura4-D18 leu1-32 | This study |
| CS332 | $h^?$ scd1-GFP::kanMX6 rga3::hphMX6 ura4-D18 leu1-32 | This study |
| CS357 | $h^-$ pact1':CRIB-3mCherry:bsdMX:ade6+ leu1-32 | This study |
| CS359 | $h^-$ pact1':CRIB-3mCherry:bsdMX:ade6+ scd1::natMX6 leu1-32 | This study |
| CS360 | $h^-$ pact1':CRIB-3mCherry:bsdMX:ade6+ scd1::natMX6 nmt81'::tea1-gef1-GFP::leu1+ leu1-32 | This study |
| CS361 | $h^+$ pact1':CRIB-3mCherry:bsdMX:ade6+ rga4-GFP::kanMX6 ura4-D18 leu1-32 | This study |
| CS362 | $h^-$ pact1':CRIB-3mCherry:bsdMX:ade6+ rga6-GFP::kanMX6 ura4-D18 leu1-32 | This study |
| CS365 | $h^-$ pact1':CRIB-3mCherry:bsdMX:ade6+ scd1::natMX6 nmt81'::tea1-scd1-GFP::leu1+ leu1-32 | This study |
| CS369 | $h^-$ pact1':CRIB-3mCherry:bsdMX:ade6+ rga6-GFP::kanMX6 sty1::natMX6 ura4-D18 leu1-32 | This study |
| CS373 | $h^-$ pact1':CRIB-3GFP:bsdMX:ade6+ scd1::natMX6 nmt81'::tea1-gef1-mCherry::leu1+ ura4-D18 leu1-32 | This study |
| CS375 | $h^-$ pact1':CRIB-3GFP:bsdMX:ade6+ gef1::kanMX6 nmt81'::tea1-gef1-mCherry::leu1+ ura4-D18 leu1-32 | This study |
| EP48 | h+ pyp1::natMX6 | This study |
| FV1218 | h- gef1-3YFP::kanMX6 leu1-32 ura4-D18 ade6-704 | (Das et al., 2015) |
| JA1329 | h- hta1-mRFP::kanMX6 | This study |
| MS98 | h- atf1::natMX6 | (Fernandez-Vazquez et al., 2013) |
| PPG56.66 | h+ scd1-GFP::kanMX6 leu1-32 ura4-D18 | This study |
| PPG65.60 | h+ ppak1::CRIB-3GFP::ura4+ ura4-D18 leu1-32 | (Revilla-Guarinos et al., 2016) |

| | | |
|---|---|---|
| PPG69.03 | *h? pak1-3GFP::kanMX6* | This study |
| PPG70.07 | *h? ppak1::CRIB-3GFP::ura4+ gef1::kanMX6 ura4-D18 leu1-32* | This study |
| PPG71.51 | *h? CRIB-3GFP::ura4+ rga4::kanMX6 ura4-D18 leu1-32* | This study |
| PPG142.42 | *h- scd2-GFP::kanMX6 leu1-32 ura4-D18* | This study |

Das, M., Nunez, I., Rodriguez, M., Wiley, D.J., Rodriguez, J., Sarkeshik, A., Yates, J.R., 3rd, Buchwald, P., and Verde, F. (2015). Phosphorylation-dependent inhibition of Cdc42 GEF Gef1 by 14-3-3 protein Rad24 spatially regulates Cdc42 GTPase activity and oscillatory dynamics during cell morphogenesis. Mol Biol Cell *26*, 3520-3534.

Fernandez-Vazquez, J., Vargas-Perez, I., Sanso, M., Buhne, K., Carmona, M., Paulo, E., Hermand, D., Rodriguez-Gabriel, M., Ayte, J., Leidel, S.*, et al.* (2013). Modification of tRNA(Lys) UUU by elongator is essential for efficient translation of stress mRNAs. PLoS Genet *9*, e1003647.

Leupold, U. (1970). Genetical methods for *Schizosaccharomyces pombe*. Methods Cell Physiol *4*, 169-177.

Revilla-Guarinos, M.T., Martin-Garcia, R., Villar-Tajadura, M.A., Estravis, M., Coll, P.M., and Perez, P. (2016). Rga6 is a Fission Yeast Rho GAP Involved in Cdc42 Regulation of Polarized Growth. Mol Biol Cell.

Zuin, A., Vivancos, A.P., Sanso, M., Takatsume, Y., Ayte, J., Inoue, Y., and Hidalgo, E. (2005). The glycolytic metabolite methylglyoxal activates Pap1 and Sty1 stress responses in Schizosaccharomyces pombe. J Biol Chem *280*, 36708-36713.

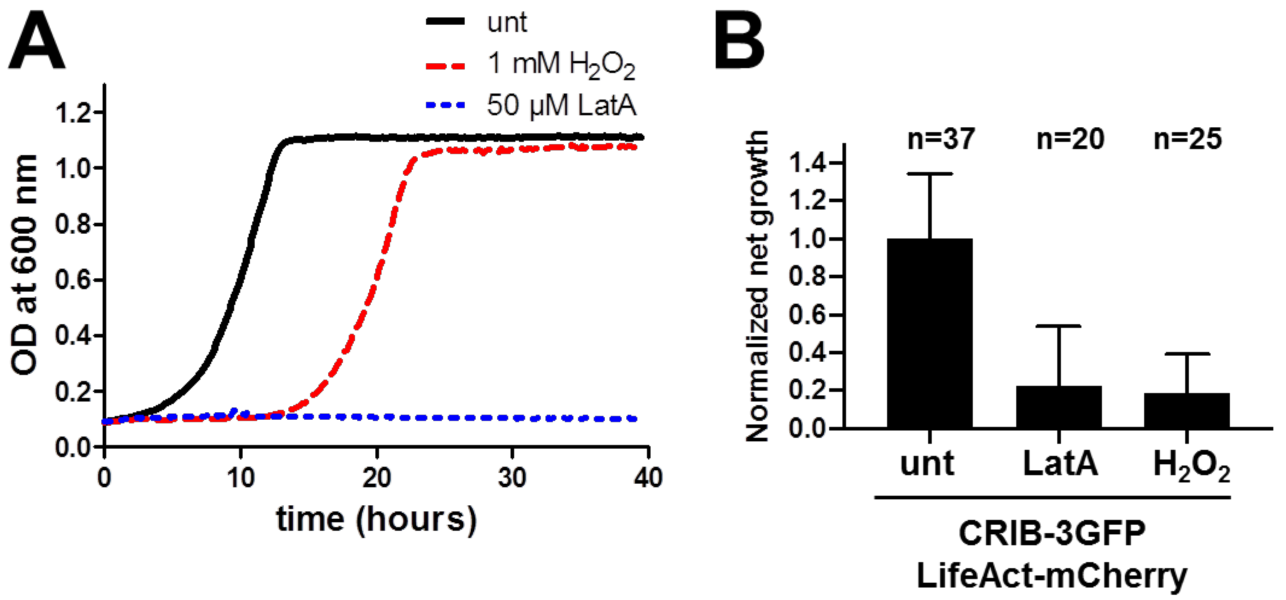


**Figure S1. Stress imposition causes a Cdc42-dependent growth arrest.**

(**A**) $H_2O_2$ causes a transient growth arrest, while LatA growth inhibition is permanent. $OD_{600nm}$ of YE cultures as in Fig 1A were treated with 1 mM $H_2O_2$, 50 μM LatA or were left untreated.

(**B**) Cell growth is inhibited after either LatA or $H_2O_2$ imposition. Net cell elongation during 48 min was calculated from movies as in Fig 1A. The measurements were normalized to the mean value of untreated condition. Means, standard deviations (SD) and number of cells analyzed (n) are shown.

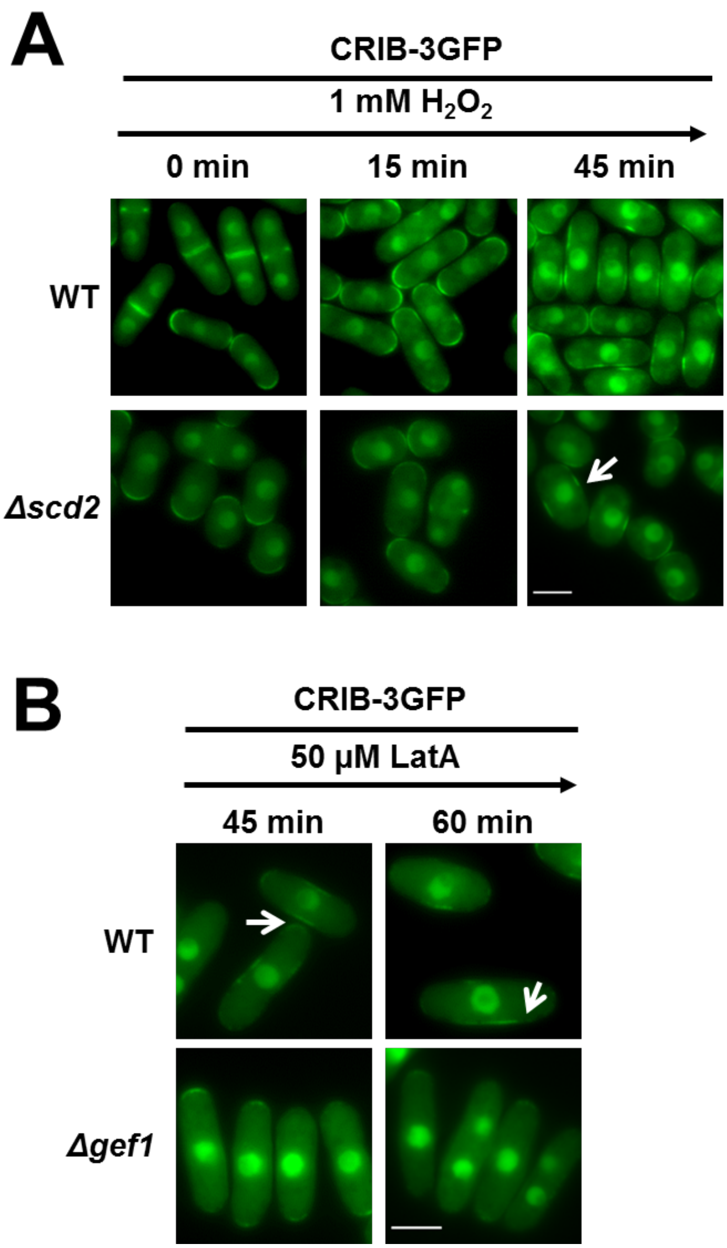


**Figure S2. Gef1, but not Scd2, is required for the formation of GTP-Cdc42 lateral patches.**

(**A**) The scaffold Scd2 is dispensable for Cdc42 activation at lateral patches. Representative images of GTP-bound Cdc42 from logarithmic YE cultures of strains PPG65.60 (*CRIB-3GFP*) and CS281 (*CRIB-3GFP Δscd2*) treated with 1 mM $H_2O_2$ for the indicated time-points. White arrows indicate the presence of Cdc42-GTP lateral patches. Scale bar: 5 µm.

(**B**) Lateral patch formation upon LatA treatment also relies on Gef1. Representative images of GTP-bound Cdc42 from logarithmic YE cultures of strains PPG65.60 (*CRIB-3GFP*) and PPG70.07 (*CRIB-3GFP Δgef1*) treated with 50 µM LatA for the indicated time-points. White arrows indicate the presence of Cdc42-GTP lateral patches. Scale bar: 5 µm.

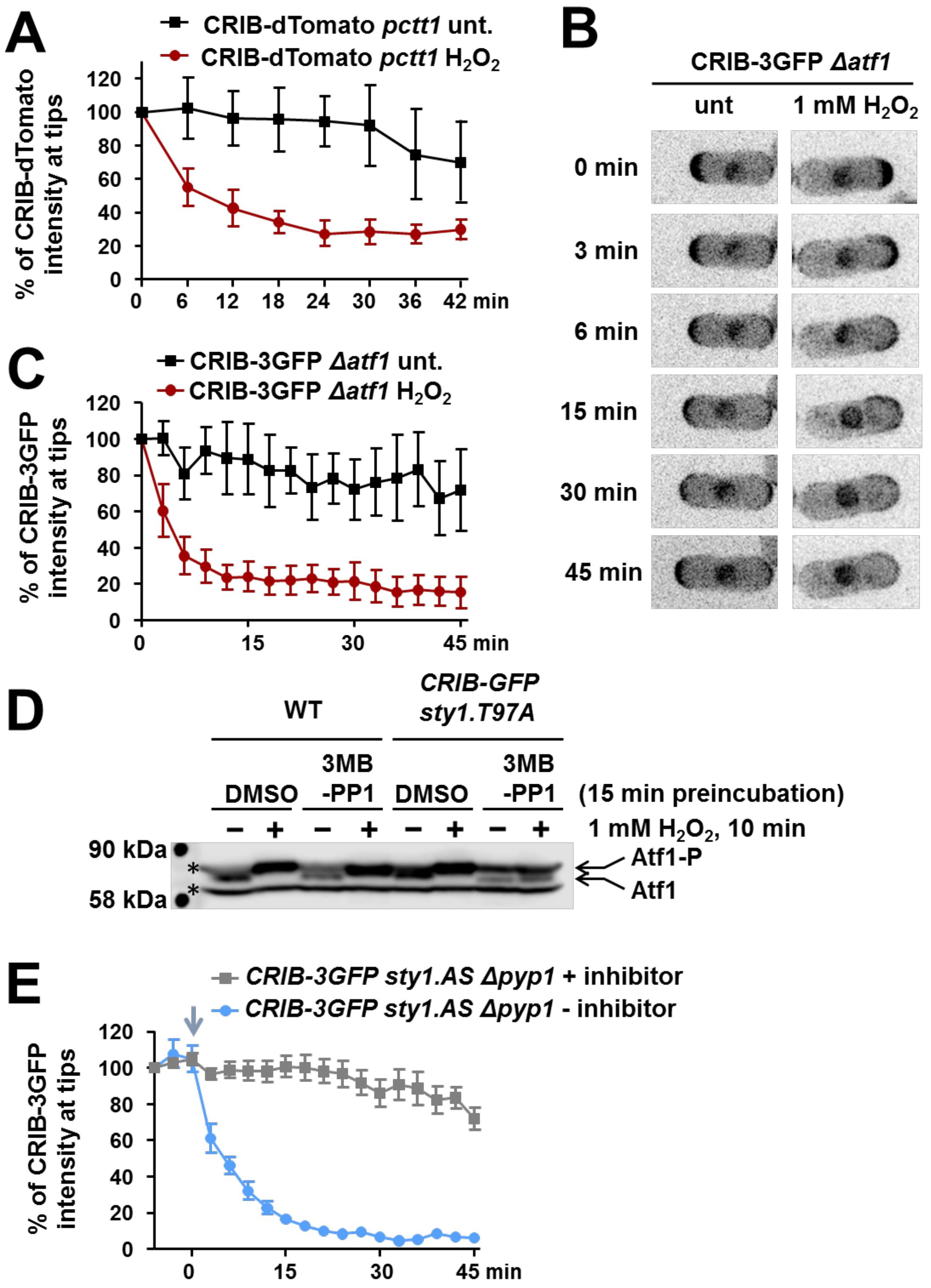


**Figure S3. GTP-Cdc42 depolarization from cell tips does not rely on gene transcription.**

(**A**) The overexpression of catalase does not delay GTP-Cdc42 depolarization after $H_2O_2$. Quantification of CRIB-dTomato intensity at cell tips from movies as in Fig. 3A. Mean and standard deviation (SD) are shown. At least ten cells were analyzed for each timepoint.

(**B**) The absence of Atf1, the main transcription factor downstream Sty1, does not affect Cdc42 depolarization from cell tips. Still images showing GTP-bound Cdc42 at different points from time-lapse experiments of YE cultures of strain CS133 (*CRIB-3GFP Δatf1*) in untreated conditions or after 1 mM $H_2O_2$ imposition.

(**C**) Quantification of CRIB-3GFP intensity at cell tips from movies as in B. Means and standard deviations (SD) are shown. At least ten cells were analyzed for each timepoint.

(**D**) Analysis of Atf1 phosphorylation after inhibition of the Sty1.AS mutant. YE cultures of 972 (WT) and CS136 (*CRIB-3GFP sty1.T97A*) were pre-incubated with 10 μM 3MB-PP1 and treated or not with 1 mM $H_2O_2$. TCA extracts were obtained and analyzed by Western blot using polyclonal antibodies (anti-Atf1).

(**E**) Quantification of CRIB-3GFP intensity at cell tips from movies as in Fig. 2F. Means and standard deviations (SD) are shown. Blue arrow indicates the time-point when the ATP-competitive inhibitor 1NM-PP1 was removed.

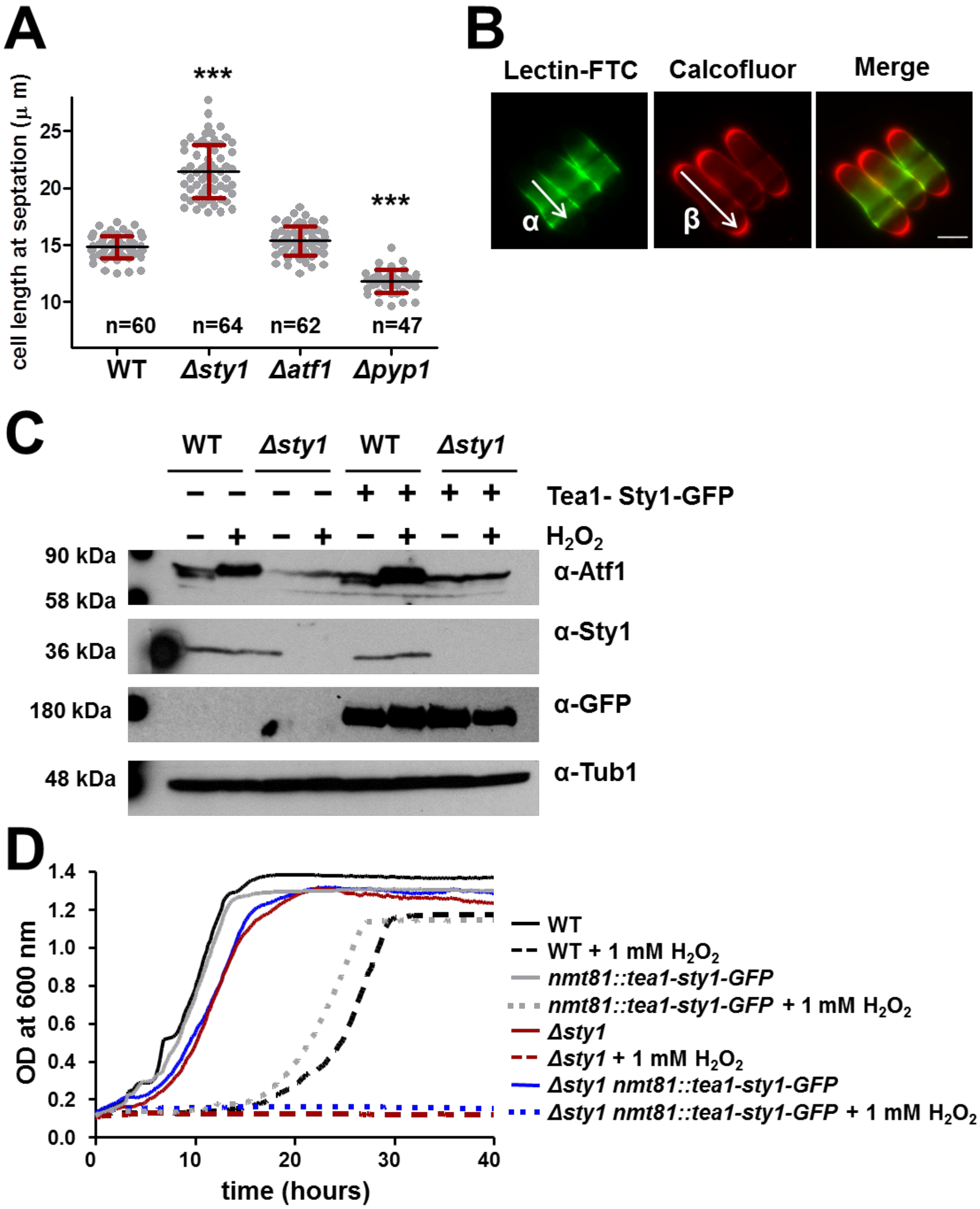


**Figure S4. Tip-anchored Sty1 kinase is not able to activate the stress-dependent transcriptional response.**

(**A**) Cell length differences are exacerbated on MM growth. Cell length distribution of 972 (WT), AV18 (*Δsty1*), MS98 (*Δatf1*) and EP48 (*Δpyp1*) on MM cultures. Mean, SD and number of cells analyzed (n) are indicated. Statistical significance was calculated using unpaired t-test (***p-value>0.0001).

(**B**) Representation of the approach used to calculate elongation rates. Lectin-FITC stains cell surface composition at the initial point. After growing for 90 min, the same culture is counterstained with calcofluor. Elongation rate is calculated as the difference between "α" and "β" divided by the growth time.

(**C**) Tip-anchored Sty1 is not able to phosphorylate the nuclear transcription factor Atf1. MM cultures of 972 (WT), AV18 (*Δsty1*), CS309 (*nmt81::tea1-sty1-GFP::leu1+*) and CS310 (*Δsty1 nmt81::tea1-sty1-GFP::leu1+*) were treated or not with 1 mM $H_2O_2$ for 15 min. TCA protein extracts were prepared and analyzed by Western blot using the indicated antibodies.

(**D**) Tip-anchored Sty1 is not able to rescue the sensitivity to $H_2O_2$ of *Δsty1* cells. Growth curves of MM cultures of strains as in C treated or not with with 1 mM $H_2O_2$ are shown.

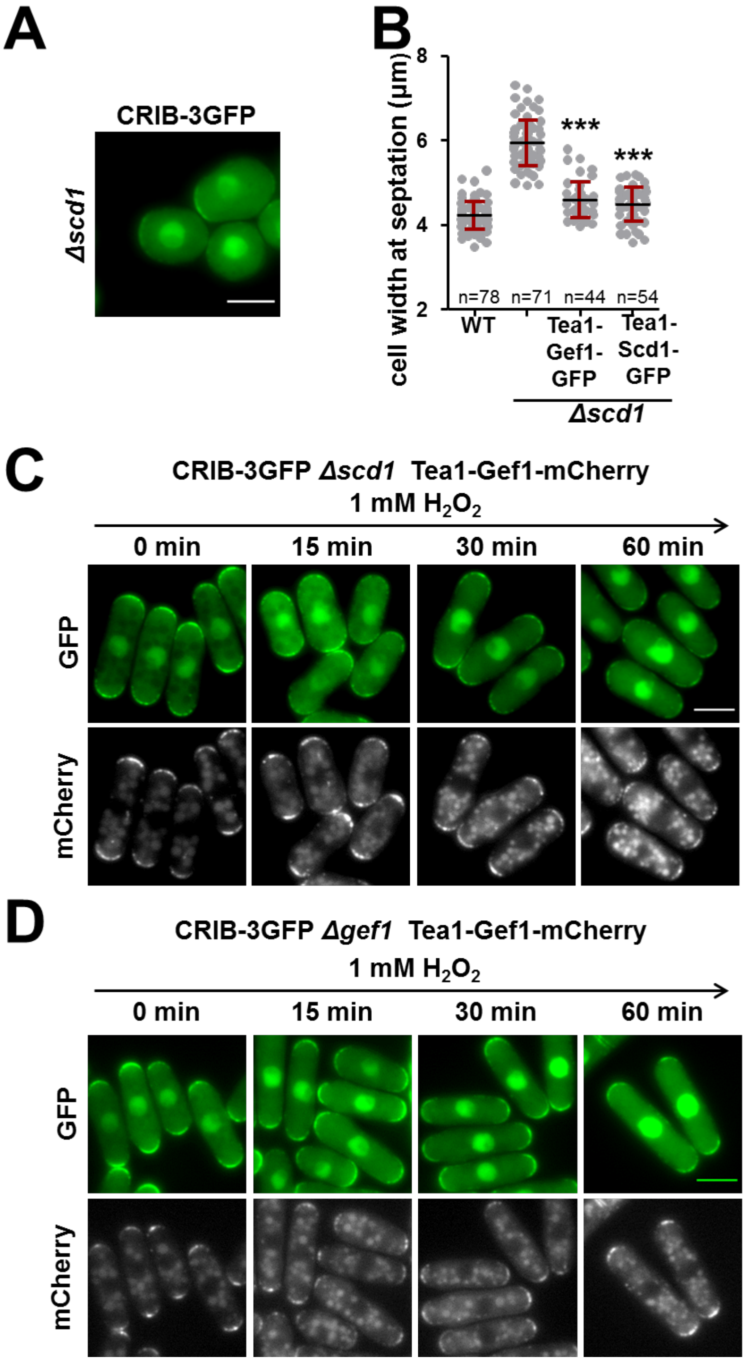


**Figure S5. Tip-anchored Gef1 is not inhibited after oxidative stress.**

(**A**) CRIB-3GFP levels are hardly observable at cells lacking *Δscd1*. Representative image showing CRIB-3GFP staining of YE culture of CS229 (*CRIB-3GFP Δscd1*). Scale bar: 5 µm.

(**B**) Trapping GEFs to cell tips restores the cell width defects of *Δscd1* strains. Cell width distribution at septation of the same strains as in Fig. 6A. Mean, SD and number of cells analyzed (n) are indicated. Statistical significance was calculated using unpaired t-test (***p-value>0.0001).

(**C-D**) Cells expressing Tea1-Gef1-mCherry are also blinded for Cdc42 depolarization to stress imposition. Representative images showing CRIB-3GFP and mCherry staining of MM cultures of CS373 (*CRIB-3GFP Δscd1 nmt81::tea1-gef1-mCherry*) and CS375 (*CRIB-3GFP Δgef1 nmt81::tea1-gef1-mCherry*) treated with 1 mM $H_2O_2$ for the indicated times. Scale bar: 5 µm.

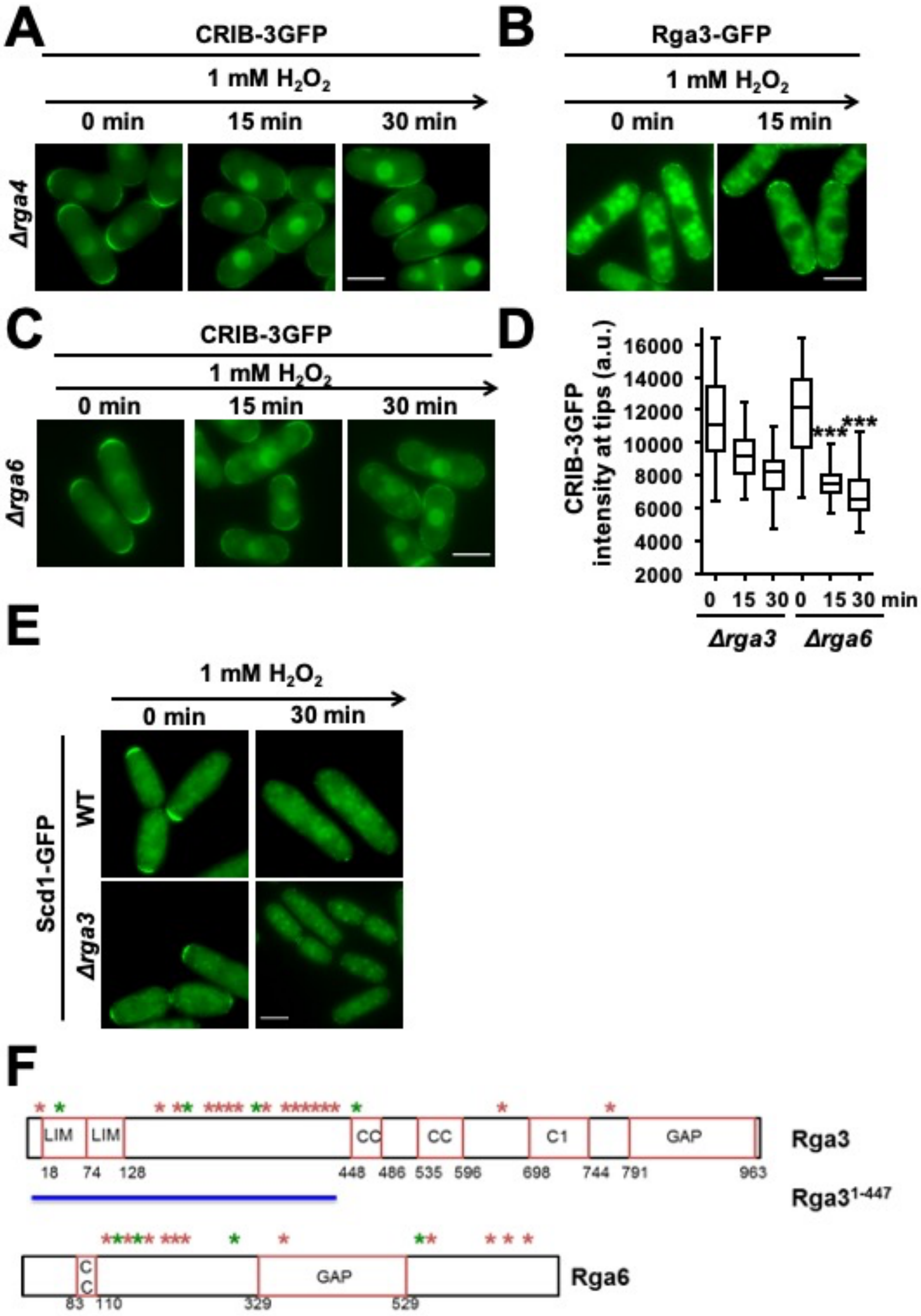


**Figure S6. Role of GAPs in stress-dependent Cdc42 inhibition.**

(**A**) Cells lacking Rga4 suffer Cdc42 depolarization after $H_2O_2$ imposition. Representative images of CRIB-GFP staining of YE cultures of PPG71.51 (*CRIB-3GFP Δrga4*) treated with 1 mM $H_2O_2$ for the indicated times. Scale bar: 5 µm.

(**B**) Rga3 remains at cell tips after $H_2O_2$ imposition. Representative images showing Rga3-GFP staining of YE culture of CS235 (*rga3-GFP*) treated with 1 mM $H_2O_2$ for the indicated times. Scale bar: 5 µm.

(**C**) The single deletion of *rga6* is not sufficient to halt Cdc42 depolarization after stress imposition. Representative images of GTP-bound Cdc42 from YE cultures of PPG75.50 (*CRIB-3GFP Δrga6*) treated with 1 mM $H_2O_2$ for the indicated time-points. Scale bar: 5 μm.

(**D**) Quantification of CRIB-3GFP intensity at cell tips of the same strains as in Fig. 5E and S6C. Representation as described in Fig. 5E. Fifty tips were analyzed for each condition within at least two independent biological replicates. The statistical significance in regard to single CRIB-3GFP *Δrga3* cells is indicated.

(**E**) Scd1 is gone from cell tips after stress imposition in *Δrga3* cells. Representative images of GFP-tagged Scd1 from MM culture of PPG56.66 (*scd1-GFP*) and CS332 (*scd1-GFP Δrga3*) treated with 1 mM $H_2O_2$ for the indicated time-points. Scale bar: 5 μm.

(**F**) Rga3 canonical S/T-P sites are located at the N-terminus of the protein. Schematic representation of Rga3 and Rga6 proteins. The different domains and the location of the canonical phosphorylation sites is indicated. Red asteriks represent S-P sites, while green asterisks mark T-P sites. LIM-:Zinc-binding domain. CC: coiled coil region C1: protein kinase C conserved region. RhoGAP: GTPase-activator protein for Rho-like GTPases.
